# E2f coordinates the proliferation and transdifferentiation of a Sox9^+^bile duct cell to initiate hepatocellular carcinoma

**DOI:** 10.1101/2025.09.18.677170

**Authors:** Gabriella Crosier, Eunsun Kim, Katharina Hayer, Emma F Furth, Patrick Viatour

## Abstract

Disruption of the Rb/E2f interaction is one of the six original hallmarks of cancer and is thought to confer unrestricted proliferative potential to tumor cells. Although E2f is active in almost all cancer cases as a result of this disruption, the consequence of its unrestricted activity for cancer initiation remains mostly unknown. To address this question, we genetically inactivated the entire *Rb* family of genes (Triple Knock Out, *TKO*) in specific cell lineages in the liver. Strikingly, this unbiased approach reveals that *Rb* family inactivation initiates a phenotypically similar hepatocellular carcinoma (HCC) from either periportal hepatocytes or *Sox9^+^* bile duct cells. In the latter population, *Rb* family inactivation results in unrestricted proliferation as well as the E2f-driven transactivation of the pioneer factor *Foxa3*, which activates a “periportal hepatocyte-like” metabolic program to ensure transdifferentiation into hepatocytes. Notably, single or compound inactivation of *E2f1* and *E2f3* in the *TKO* background is sufficient to suppress *Foxa3* expression and HCC initiation, without altering the proliferative status of Sox9^+^ bile duct cells. Collectively, our results reveal that derepression of a cell cycle gene program and activation of non-cell cycle oncogenic features by E2f are distinct consequences of disrupting the Rb/E2f interaction, and that coordinated activation of these two functions are essential for HCC initiation from a founder liver bile duct cell lineage.

## INTRODUCTION

Since their identification in the late 80’s, the role of *E2f* family of transcription factors (activator *E2f1-3* and repressor *E2f4-8*) in cell cycle and the regulation of their activity by Rb family proteins (*Rb*, *p107* and *p130*) has been abundantly documented^1–3^. The disruption of the inhibitory interaction between Rb and E2f has been identified as one of the six original hallmarks of cancer^4^, predominantly on the basis of *in vitro* studies and clinical observations. This disruption is an early event in tumor initiation and the resulting unrestricted E2f activity is commonly thought to confer “insensitivity to antigrowth signals” via the activation of a cell cycle gene program. Clinical effort to target the consequences of disrupted Rb/E2f interaction has led to the successful development of CDK inhibitors^5^. Unfortunately, their limited efficacy as well as the emergence of therapeutic resistance begs for the development of alternative approaches to repress the consequences of the disruption of the Rb/E2f interaction for novel cancer therapy^6,7^. Although E2f binds to more than 10,000 genomic loci in cancer cells^8^, our understanding of E2f oncogenic activity is mostly restricted to its “canonical” role in cell cycle regulation. Therefore, whether unrestricted E2f activates “non-canonical” features that are critical to initiate tumorigenesis, and whether these non-canonical features represent novel opportunities to target oncogenic E2f activity, represents a critical question in basic and translational cancer biology.

Hepatocellular carcinoma (HCC) is the most frequent form of liver cancer^9–11^ and its main etiological factors affect both hepatocytes and bile duct cells. Hepatocytes are a *bone fide* cell of origin for HCC but the contribution of bile duct cells to HCC initiation is still debated^12^. In particular, although hepatocytes can act as a cell of origin for cholangiocarcinoma via transdifferentiation^13^, whether bile duct cells display similar plasticity in the context of HCC initiation has never been reported.

Disruption of the Rb/E2f interaction is a common consequence of the most frequent HCC initiating events (such as methylation of the *p15*/*p16* promoter, overexpression of *Cyclind1*, expression of viral oncoproteins, etc)^14–16^, suggesting that unrestricted E2f activity is critical to HCC initiation. In support of this model, we previously showed that liver-specific inactivation of *Rb* family genes (*Rb^lox/lox^*, *p130^lox/lox^*, *p107^-/^*^-^; conditional Triple Knock Out; *cTKO*) is sufficient to initiate HCC that reflects many clinical features of the human disease^14,17,18^. Interestingly, *cTKO*-initiated HCC originates from the periportal region, an area of the liver that hosts both periportal hepatocytes and bile duct cells. We also previously found that unrestricted E2f activity progressively expands its transcriptome to include a set of low-binding affinity target genes constituting a non-canonical function of E2f that encompasses features such as Notch signaling and the Warburg effect ^18–20^. However, whether the activation of these non-canonical features by E2f is required for *cTKO*-initiated HCC is unknown.

To address these questions, we developed a mouse genetic screen to probe the consequences of disrupting the Rb/E2f interaction in distinct liver cell subpopulations. This approach reveals that unrestricted E2f activity initiates a phenotypically similar HCC from both periportal hepatocytes and Sox9^+^ bile duct cells. Mechanistically, we find that E2f activates the expression of the pioneer factor *Foxa3* in Sox9^+^ bile duct cells, triggering a “periportal-like” metabolic activity that rewires their cellular identity to the hepatocyte lineage. In addition, we show that deficiency of activator *E2f1* or *E2f3* in the *cTKO* background is sufficient to inhibit *Foxa3* transactivation and prevent the malignant transformation of Sox9^+^ bile duct cells without repressing their ectopic proliferation, resulting instead in the development of benign bile duct tumors. Our findings prompt further examinations of the cellular origin of HCC, with possible implications for improving its therapeutic management.

## RESULTS

### A genetic screen to determine the cell of origin for HCC initiated upon Rb family inactivation

Pan expression of the *Cre* recombinase (intrasplenic injection of Adeno-*Cre*) in *cTKO* liver simultaneously initiates HCC lesions in all liver lobes as well as several benign cell-cycle related phenotypes, such as increased polyploidy and mitotic figures in hepatocytes and hamartoma (a benign bile duct growth) (**Fig.1a**). This compound phenotype suggests that *Rb* family inactivation impacts both hepatocytes and bile duct cells, although the outcome varies based on the cellular context.

**Figure 1.**
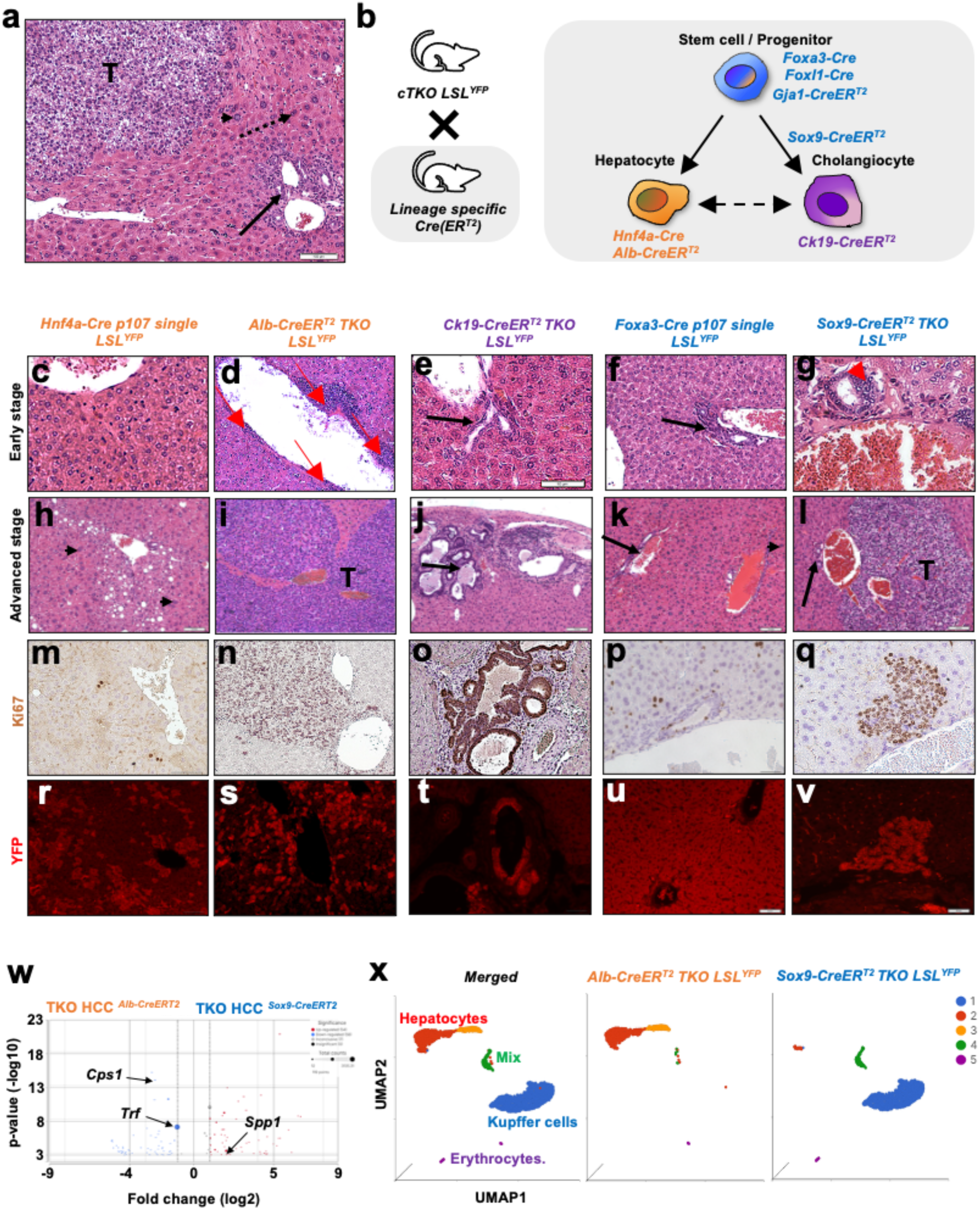
A genetic screen to identify the cell of origin for cTKO HCC. **a:** Benign and malignant phenotypes simultaneously develop in *cTKO* mice upon injection of Ad-Cre: HCC (T), increased frequency of polyploid hepatocytes (arrowhead), increased frequency of mitotic figures (dotted arrow) and hamartoma (long arrow). **b:** Experimental lineage tracing strategy to identify the cell of origin for TKO HCC. **c**-**v:** Immuno-histological analysis of representative livers from the different transgenic lines of interest at early and advanced stages. **c**-**l**: H&E staining of liver from *Hnf4a-Cre p107 single LSL^YFP^*, *Alb-CreER^T^*^2^ *TKO LSL^YFP^*, *Ck19Cre-ER^T^*^2^ *TKO LSL^YFP^*, *Foxa3-Cre p107 single LSL^YFP^* and *Sox9-CreER^T^*^2^ *TKO LSL^YFP^*, respectively at early time points (3-5 weeks, **c**-**g**), and advanced time points (6-8 weeks for *Alb-CreER^T^*^2^ *TKO LSL^YFP^*, 20-25 weeks for the other lines, either after birth (*Hnf4a-Cre* and *Foxa3-Cre*) or Tamoxifen treatment (*Ck19-CreER^T^*^2^ and *Sox9-CreER^T^*^2^), **h**-**l**). **m**-**q**: Representative Ki67 IHC staining at advanced stages. **r**-**v**: Representative immunofluorescence (IF) staining for YFP in the livers at advanced stages. **w**: Volcano plot display of comparative bulk RNA-Seq analysis for tumor cells from *Sox9-CreER^T^*^2^ *TKO LSL^YFP^* (TKO HCC *^Sox^*^9*-CreERT*2^-20-week time point) and *Alb-CreER^T^*^2^ *TKO LSL^YFP^*(TKO HCC *^Alb-CreERT^*^2^-8 week-time point). **x**: UMAP display of scRNA-Seq datasets from YFP^+^ cells isolated from either *Alb-CreER^T^*^2^ *TKO LSL^YFP^* (6-week time point) or *Sox9-CreER^T^*^2^ *TKO LSL^YFP^*(9-week time point). Left: merge display; cluster are identified based on marker expression); Center and Right: individual displays.

To identify the cellular and molecular mechanism of HCC initiation upon Rb family inactivation, we generated a series of *cTKO* mouse strains expressing either *Cre* or tamoxifen-inducible *Cre-ER^T^*^2^ recombinases from the endogenous loci of genes that are markers for distinct cell populations in the liver (hepatocytes, *Hnf4a-Cre*^21^*, Albumin-CreER^T^*^2^ ; cholangiocytes: *Ck19-CreER^T^*^2^ ^22^; various progenitors populations: *Foxa3-Cre* ^23^, *Foxl1-Cre* ^24^, *Sox9-CreER^T^*^2^ ^25^ and *Gja1-CreER^T^*^2^ ^26^) (**Fig.1b**). We further crossed these mice with a *Rosa26-LSL^YFP/+^* reporter strain that enables permanent labeling for lineage tracing purposes. Several of these mouse lines display one or more benign cell-cycle-related phenotype(s), as originally observed in Ad-*Cre* injected *cTKO* mice: polyploidy in *Hnf4a-Cre p107single (Rb^lox/lox^, p130^lox/lox^, p107^+/-^) LSL^YFP^* and *Hnf4a-Cre TKO LSL^YFP^* mice (**Fig.1c, h, m and r; Fig.S1a**), hamartoma in *Ck19-CreER^T^*^2^ *TKO LSL^YFP^* mice (**Fig.1e, j, o and t**), hepatocyte polyploidy and hamartoma in *Foxa3-Cre p107single LSL^YFP^* and *Foxa3-Cre TKO LSL^YFP^* mice (**Fig.1f, k, p and u; Fig.S1b**) and increased mitotic figure in *Foxl1-Cre TKO* mice (**Fig.S1c**). *Gja1-CreER^T^*^2^ *TKO* mice exhibit no abnormal phenotype in the liver (**Fig.S1d**). In contrast, both *Alb-CreER^T^*^2^ *TKO LSL^YFP^* and *Sox9-CreER^T^*^2^ *TKO LSL^YFP^* mice develop HCC that is histologically identical to the Adeno-*Cre*-induced *cTKO* HCC (**Fig.1d, i, n and s; Fig.1g, l, q and v; Fig.S1e**), albeit the kinetics of HCC development drastically differ between both lines (macroscopic tumors visible after 6-8 weeks and 25 weeks for TKO HCC *^Alb-CreERT^*^2^ and TKO HCC *^Sox^*^9^*^-CreERT^*^2^, respectively). In parallel to HCC, *Sox9-CreER^T^*^2^ *TKO LSL^YFP^* mice also develop hamartoma, showing that these mice simultaneously develop a dual phenotype composed of benign (hamartoma) and malignant (HCC) growths.

Comparative bulk RNA-Seq analysis of TKO HCC *^Alb-CreERT^*^2^ and TKO HCC *^Sox^*^9^*^-CreERT^*^2^ harvested at the “advanced stage” time points shows limited transcriptional differences between them (112 genes differentially regulated in total-**Fig.1w; Table S1**), confirming that these tumors are essentially similar. In contrast, comparative scRNA-Seq analysis of YFP^+^ cells (see Methods for experimental strategy) isolated at early time points (4 weeks for TKO HCC *^Alb-CreERT^*^2^ and 9 weeks for TKO HCC *^Sox^*^9^*^-CreERT^*^2^) from both models reveals that, while early tumor cells from *Alb-CreER^T^*^2^ *TKO LSL^YFP^* mice overwhelmingly display a “Hepatocyte” signature, early tumor cells from *Sox9-CreER^T^*^2^ *TKO LSL^YFP^* mice are partitioned in two groups with either “Hepatocyte” or “Mix” (expressing bile duct markers as well as some hepatocyte markers such as *Krt18*/*Krt8*) signatures, respectively, suggesting a distinct cellular origin for TKO HCC *^Alb-CreERT^*^2^ and TKO HCC *^Sox^*^9^*^-CreERT^*^2^ (**Fig.1x**; **Fig.S1g; Table S2**). Finally, analysis of Ki67 expression at the single cell level suggests lower proliferative activity in this “Mix” population (**Fig.S1h**). Collectively, these results show that inactivation of *Rb* family members in either *Albumin*- or *Sox9*-expressing cells initiates phenotypically similar HCC lesions.

### Rb family inactivation initiates HCC from both periportal hepatocytes and Sox9^+^ bile duct cells

In adult mouse liver, albumin is a pan-hepatocyte marker, while *Sox9* expression is restricted to a subset of ductular progenitors (*Sox9^high^*) and hybrid periportal hepatocytes (*Sox9^low^*)^25,27^ ^28^. Although the latter population is not thought to serve as the cell of origin in mouse models of HCC induced by carcinogens such as dimethylnitrosamine (DEN) or carbon tetrachloride (CCl4)^24,28,29^, genetic models of HCC have suggested a role for them in HCC initiation^12^. In contrast, whether *Sox9^+^* bile duct cells can serve as a cell of origin for HCC remains unresolved.

To identify the cell of origin for TKO HCC *^Alb-CreERT^*^2^ and TKO HCC *^Sox^*^9^*^-CreERT^*^2^, we first analyzed proliferation at early time points in both models by assessing Ki67 (a general marker of proliferation) expression by IHC. In *Sox9-CreER^T^*^2^ *TKO LSL^YFP^* mice, proliferation at the one week-time point is restricted to cells lining the bile ducts (**Fig.2a**). The bile duct architecture is progressively disrupted and both hamartoma and small HCC lesions can be observed by nine weeks, confirming that the inactivation of *Rb* family in *Sox9^+^* bile duct cells initiates a dual phenotype characterized by both malignant (i.e. HCC) and benign (i.e. hamartoma) growths. In contrast, proliferation one week after *Rb* family inactivation is restricted to periportal hepatocytes in *Alb-CreER^T^*^2^ *TKO LSL^YFP^* mice. While proliferation progressively expands to the entire liver, bile ducts remain quiescent (**Fig.2b**), confirming the distinct cellular origins of TKO HCC *^Alb-^_CreERT2_* versus TKO HCC *_Sox9-CreERT2_*.

**Figure 2.**
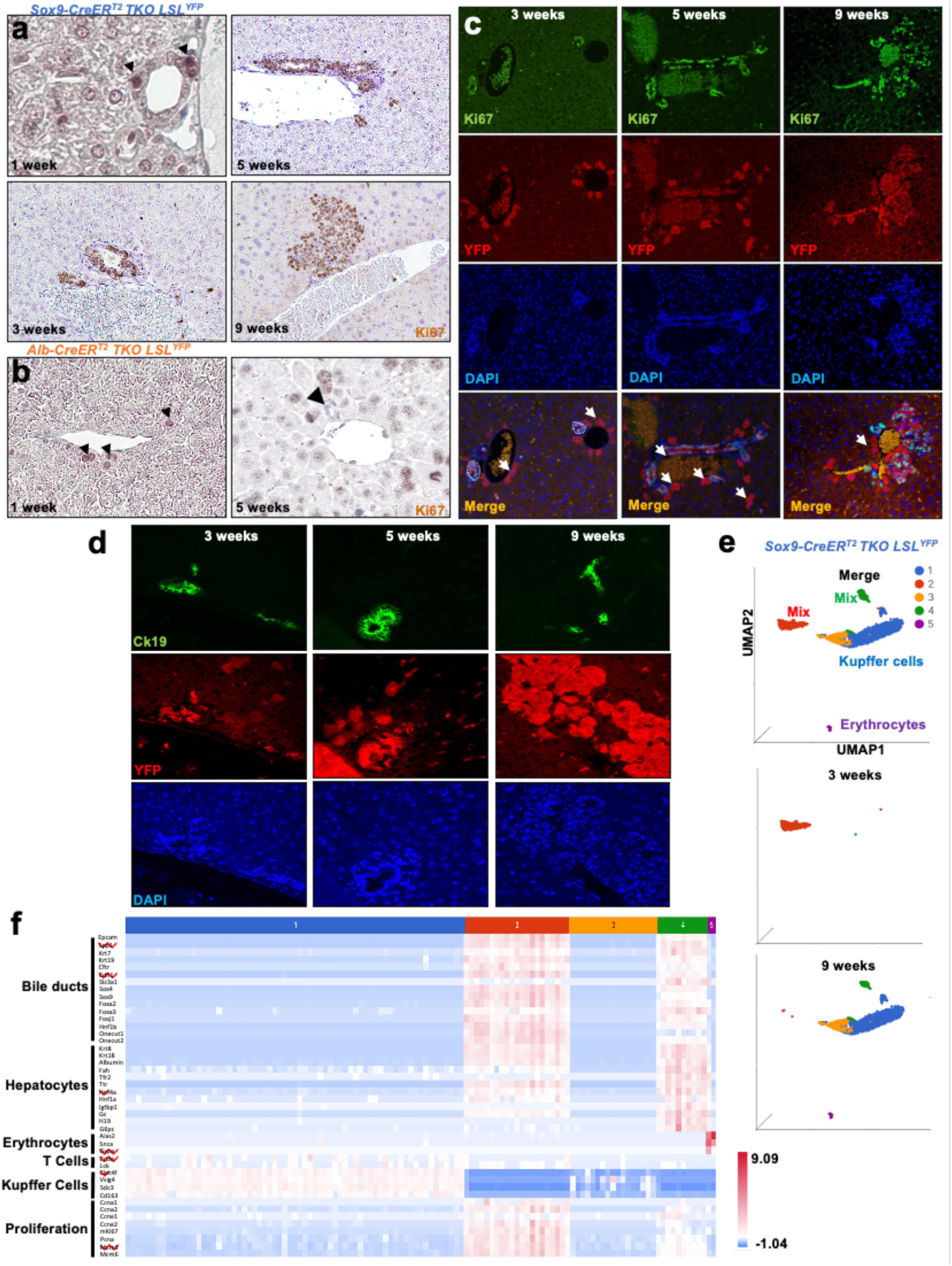
TKO HCC *^Sox^*^9*-CreERT*2^ originates from bile ducts. **a**: Representative Ki67 IHC staining of livers from *Sox9-CreER^T^*^2^ *TKO LSL^YFP^* mice at one-(arrowheads indicate proliferative bile duct cells), three-, five-, and nine-weeks post Tamoxifen treatment. **b**: Representative Ki67 IHC staining of livers from *Alb-CreER^T^*^2^ *TKO LSL^YFP^* mice at one week (arrowheads indicate proliferative periportal hepatocytes) and five weeks (arrowhead indicates non-proliferative bile duct) post Tamoxifen treatment. **c**: Co-IF staining of livers from *Sox9-CreER^T^*^2^ *TKO LSL^YFP^* mice at three-, five- and nine-weeks post Tamoxifen treatment for Ki67 and YFP counterstained with DAPI, displayed either individually or merged (bottom row). Dotted circles indicate bile ducts; arrowheads indicate recombined hepatocytes. **d**: Co-immunostaining of livers from *Sox9-CreER^T^*^2^ *TKO LSL^YFP^* mice at three-, five- and nine-weeks post Tamoxifen treatment for Ck19 and YFP counterstained with DAPI, displayed individually. **e**: UMAP display of merged scRNA-Seq datasets from YFP^+^ cells isolated from *Sox9-CreER^T^*^2^ *TKO LSL^YFP^* liver at three- and nine-week time points after Tamoxifen treatment. Upper panel: merge display; middle and lower panels: individual displays. **f**: heatmap showing key lineage and proliferation marker expression in merged scSeq-RNA datasets from *Sox9-CreER^T^*^2^ *TKO LSL^YFP^* at three- and nine-week time points, as displayed in **Fig.2e.**

To further characterize the ductal origin of TKO HCC *^Sox^*^9^*^-CreERT^*^2^, we performed lineage tracing analysis by immunofluorescence (IF) detection of YFP expression in *Sox9-CreER^T^*^2^ *TKO LSL^YFP^* liver. This analysis confirms allelic recombination in both bile duct and periportal cells (**Fig.2c; Fig.S2a**). However, only YFP^+^ bile duct cells are positive for Ki67 (**Fig.2c**) and BrDU (**Fig.S2a**), confirming that proliferative activity is restricted to *Sox9^+^* bile duct cells upon *Rb* family inactivation. Next, we performed a similar lineage tracing analysis by combining the detection of YFP with lineage markers Ck19 (bile duct) and Hnf4a (hepatocyte) in *Sox9-CreER^T^*^2^ *TKO LSL^YFP^* liver (**Fig.2d**; **Fig.S2b**). YFP^+^ cells initially express Ck19 but progressively lose its expression. In contrast, YFP^+^ cells do not initially express Hnf4a but progressively gain its expression, suggesting the progressive transdifferentiation of Sox9^+^ bile duct cells towards the hepatocyte lineage upon *Rb* family inactivation. To characterize this progressive transdifferentiation at the molecular level, we performed a comparative scRNA-Seq analysis of YFP^+^ cells isolated from *Sox9-CreER^T^*^2^ *TKO LSL^YFP^* mice at either the three- or nine-week time points. Although cells from both time points express a “Mix” signature (simultaneous expression of bile duct and hepatocyte markers), they cluster separately (**Fig.2e; Table S3**). Beyond different proliferative rates (**Fig. S2c**), analysis of lineage marker expression shows that tumor cells at the three-week time point express higher levels of bile duct markers and lower level of hepatocyte markers compared to their nine-week time point counterpart (**Fig.2f**; **Fig.S2d-f**), confirming the progressive transdifferentiation of Sox9^+^ bile duct cells towards the hepatocyte lineage upon *Rb* family inactivation.

### Epcam as a marker for the prospective characterization of TKO HCC ecosystem

scRNA-Seq analysis of early time points TKO HCC *^Alb-CreERT^*^2^ and TKO HCC *^Sox^*^9^*^-CreERT^*^2^ cells (**Fig.S3a**) suggests that *Epcam* (a marker expressed at the surface of bile duct cells^30^) is differentially expressed between the “Hepatocyte” and “Mix” cluster cells. Flow cytometry analysis confirms that Epcam surface expression partitions TKO HCC *^Alb-CreERT^*^2^ and TKO HCC *^Sox^*^9^*^-CreERT^*^2^ cells into Epcam^high^ and Epcam^-^ populations: while YFP^+^ cells are exclusively Epcam^high^ in control *Sox9-CreER^T^*^2^ *LSL^YFP^* mice, Epcam^-^ cells progressively become dominant during TKO HCC *^Sox^*^9^*^-CreERT^*^2^ development to ultimately recapitulate the Epcam expression profile of TKO HCC *^Alb-CreERT^*^2^ tumor cells (**Fig3a**). IF staining for Epcam and DAPI confirms that most YFP^+^ cells express Epcam at early time points while Epcam expression becomes rare in TKO HCC *^Sox^*^9^*^-CreERT^*^2^ lesions at advanced time points (**Fig.S3b**). Comparative bulk RNA-Seq analysis of Epcam^high^ and Epcam^-^ tumor cells recapitulates the differential gene expression profile observed between “Hepatocyte lineage” and “Mix” (expression of bile duct cell markers and some hepatocyte markers such as *Krt8*/*Krt18*) clusters (**Fig.3b**; **Fig.S1f** and **Fig.S3c; Table S4**). Collectively, these data show Epcam surface marker expression correlates with its mRNA expression and enables the prospective isolation of “Hepatocyte lineage”/Epcam^-^ and “Mix”/Epcam^high^ populations for further functional and molecular characterization.

**Figure 3.**
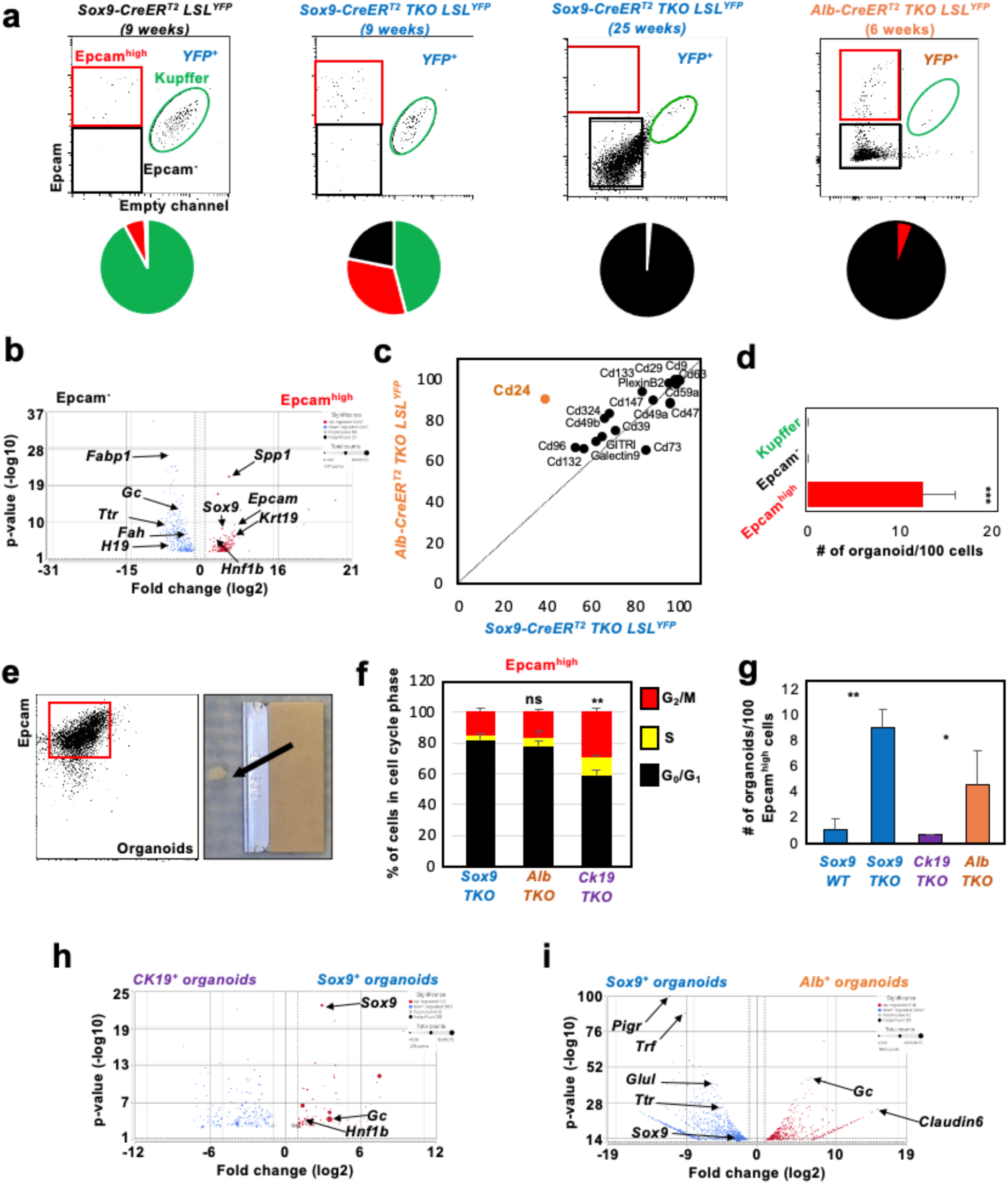
Epcam expression identifies a cellular hierarchy in TKO HCC. **a:** Identification of Kupffer cells (green), Epcam^high^ (red) and Epcam^-^ (black) cells based on Epcam expression by flow analysis. Upper panels: Representative FACS plot. Lower panels: Pie chart for the distribution of YFP^+^ cells in each group for different lines (n=5). *NB*: bulk RNA-Seq of flow isolated Kupffer cells confirm their expression of Kupffer cell specific markers (not shown). **b**: Volcano plot display of comparative bulk RNA-Seq analysis for Epcam^high^ and Epcam^-^ cells isolated from *Sox9-CreER^T^*^2^ *TKO LSL^YFP^* nine weeks post Tamoxifen treatment. The expression of key lineage markers is highlighted. **c**: Validation by flow analysis of 18 surface marker expression in Epcam^high^ cells from *Sox9-CreER^T^*^2^ *TKO LSL^YFP^* and *Alb-CreER^T^*^2^ *TKO LSL^YFP^* (n=3 per line). **d:** Organoid forming assay on 100 sorted Kupffer cells (negative control), Epcam^high^ and Epcam^-^ cells. Cells were cultured for two weeks prior to organoid counting (n=5). **e**: Left: Representative flow analysis for Epcam expression on Epcam^high^-derived organoid cells after 100 days in culture (n=3). Right: Representative subcutaneous tumor grown upon injection of organoids mixed with Matrigel (n=2/4). **f**: Cell cycle analysis (PI incorporation) of Epcam^high^ cells isolated from different lines at nine-week (*Sox9-CreER^T^*^2^ *TKO LSL^YFP^ - Sox9 TKO* and *Ck19-CreER^T^*^2^ *TKO LSL^YFP^ – Ck19 TKO*) and six week- (*Alb-CreER^T^*^2^ *TKO LSL^YFP^* -*Alb TKO*) time points (n=5). **g**: Organoid forming assay with 100 Epcam^high^ cells from *Sox9-CreER^T^*^2^ *LSL^YFP^* (*Sox9 WT*), *Sox9-CreER^T^*^2^ *TKO LSL^YFP^* (*Sox9 TKO*), *Ck19-CreER^T^*^2^ *TKO LSL^YFP^* (*Ck19 TKO*) and *Alb-CreER^T^*^2^ *TKO LSL^YFP^* (*Alb TKO*) (n=5). **h**-**i**: Volcano plot display of comparative bulk RNA-Seq analysis for organoids derived from Epcam^high^ cells from *Ck19-CreER^T^*^2^ *TKO LSL^YFP^* (*Ck19*^+^ organoids) versus *Sox9-CreER^T^*^2^ *TKO LSL^YFP^*(*Sox9*^+^ organoids) (**h**) *and Sox9-CreER^T^*^2^ *TKO LSL^YFP^* (*Sox9^+^* organoids) versus *Alb-CreER^T^*^2^ *TKO LSL^YFP^* (*Alb*^+^ organoids) (**i**).

Screening for 269 surface marker expression identifies 18 markers expressed at the surface of Epcam^high^ cells in *Sox9-CreER^T^*^2^ *TKO LSL^YFP^* mice. Among these markers, only Cd24 displays different expression between Epcam^high^ cells from *Sox9-CreER^T^*^2^ *TKO LSL^YFP^* and *Alb-CreER^T^*^2^ *TKO LSL^YFP^* mice (**Fig.3c**). However, IHC for Cd24 does not identify differential expression between TKO HCC *^Alb-CreERT^*^2^ and TKO HCC *^Sox^*^9^*^-CreERT^*^2^ cells (**Fig. S3d**), further supporting their overlapping identity at the advanced time point.

Within the TKO HCC *^Sox^*^9^*^-CreERT^*^2^ ecosystem, the organoid forming activity is restricted to the Epcam^high^ cell fraction (**Fig.3d**). Organoids derived from these cells (TKO organoids) maintain high Epcam expression level during long-term culture and form tumors upon transplantation (**Fig.3e**). Cell cycle analysis does not show increased *in vivo* proliferative activity of Epcam^high^ cells freshly isolated from *Sox9-CreER^T^*^2^ *LSL^YFP^* mice compared to their counterparts from *Alb-CreER^T^*^2^ *TKO LSL^YFP^* and *Ck19-CreER^T^*^2^ *TKO LSL^YFP^* mice (**Fig.3f**). However, they have higher organoid forming capacity (**Fig.3g**), suggesting a lack of correlation between the *in vivo* cell cycle activity and the *in vitro* growth capacity of Epcam^high^ cells from the different lines. In addition, organoid cells derived from *Sox9-CreER^T^*^2^ *TKO LSL^YFP^* mice also show higher proliferation rate compared to their *Alb-CreER^T^*^2^ *TKO LSL^YFP^* and *Ck19-CreER^T^*^2^ *TKO LSL^YFP^* counterparts (**Fig.S3e**). Of note, the observed organoid forming capacity of Epcam^high^ cells isolated from *Ck19-CreER^T^*^2^ *TKO LSL^YFP^*mice suggest a potential HCC initiating activity in these cells. Indeed, careful examination of liver sections from 15 *Ck19-CreER^T^*^2^ *TKO LSL^YFP^* mice identifies the presence of a single tumor, confirming the rare HCC initiating capacity of these cells (**Fig.S3f**). Finally, bulk RNA-Seq analysis of organoid cells derived from these different lines showed higher transcriptional variations between Sox9^+^-versus Alb^+^-derived organoids (1693 differentially regulated genes) compared to Sox9^+^-versus Ck19^+^-derived organoids (270 differentially regulated genes) (**Fig.3h-I; Table S5-6**), suggesting the higher transcriptional similarity of these latter two lines. Collectively, these data establish that Epcam expression distinguishes a cellular hierarchy within HCC initiated from *cTKO* mice.

### Activation of proliferation is not sufficient to initiate TKO HCC

Among activator *E2f* genes, *E2f1* and *E2f3* are thought to play a key role in HCC pathogenesis since *E2f2* is not expressed in HCC^31^. To determine the individual and compound role of activator *E2fs* in TKO HCC development, we crossed *Sox9-CreER^T^*^2^ *TKO LSL^YFP^* mice with *E2f1^-/-^ E2f3^lox/lox^* mice^32^ (**Fig.4a**). Individual *E2f1* and *E2f3* deficiency does not suppress the formation of benign hamartoma but largely prevents the development of HCC whereas dual *E2f1*/*E2f3* deficiency completely prevents both (**Fig.4b**; **Fig.S4a-b**). IHC staining for Ki67 (**Fig.4c**) and cell cycle analysis of Epcam^high^ cells (**Fig.4d**) show that single and dual *E2f1* and *E2f3* deficiency does not repress proliferation upon *Rb* family inactivation in Sox9^+^ bile duct cells. Interestingly, the rare TKO HCC lesions developing in a *E2f1* deficient background display a similar proliferative index compared to TKO HCC developing in a wild type *E2f* background (**Fig.S4c**). In addition, *E2f1* deficiency does not impair the organoid forming capacity of Epcam^high^ cells compared to their parental TKO counterparts (**Fig.4e**), suggesting that *E2f3* is more important than *E2f1* for TKO HCC initiation. However, both *E2f1* and *E2f3* deficiency decreases the proliferation of organoid cells (**Fig.4f**). Finally, long term (12 months) assessment of *Sox9-CreER^T^*^2^ *TKO LSL^YFP^ E2f1^-/-^ E2f3^lox/lox^* mice shows that TKO HCC does not develop in the absence of activator *E2fs* (**Fig.S4d-e**), confirming that unrestricted E2f activity is the only driver of tumorigenesis upon functional inactivation of the *Rb* family proteins. Collectively, these experiments show that *E2f1* and *E2f3* are not required *stricto sensu* for the activation of proliferation upon disruption of the Rb/E2f interaction. However, whereas the presence of either *E2f1* or *E2f3* is sufficient for the initiation of benign hamartoma development, the presence of both activator *E2f*s is required for HCC initiation. Overall, these findings strongly suggest that E2f relies upon its critical activation of non-canonical functions to transform Sox9^+^ bile duct cells into a cell of origin for HCC.

**Figure 4.**
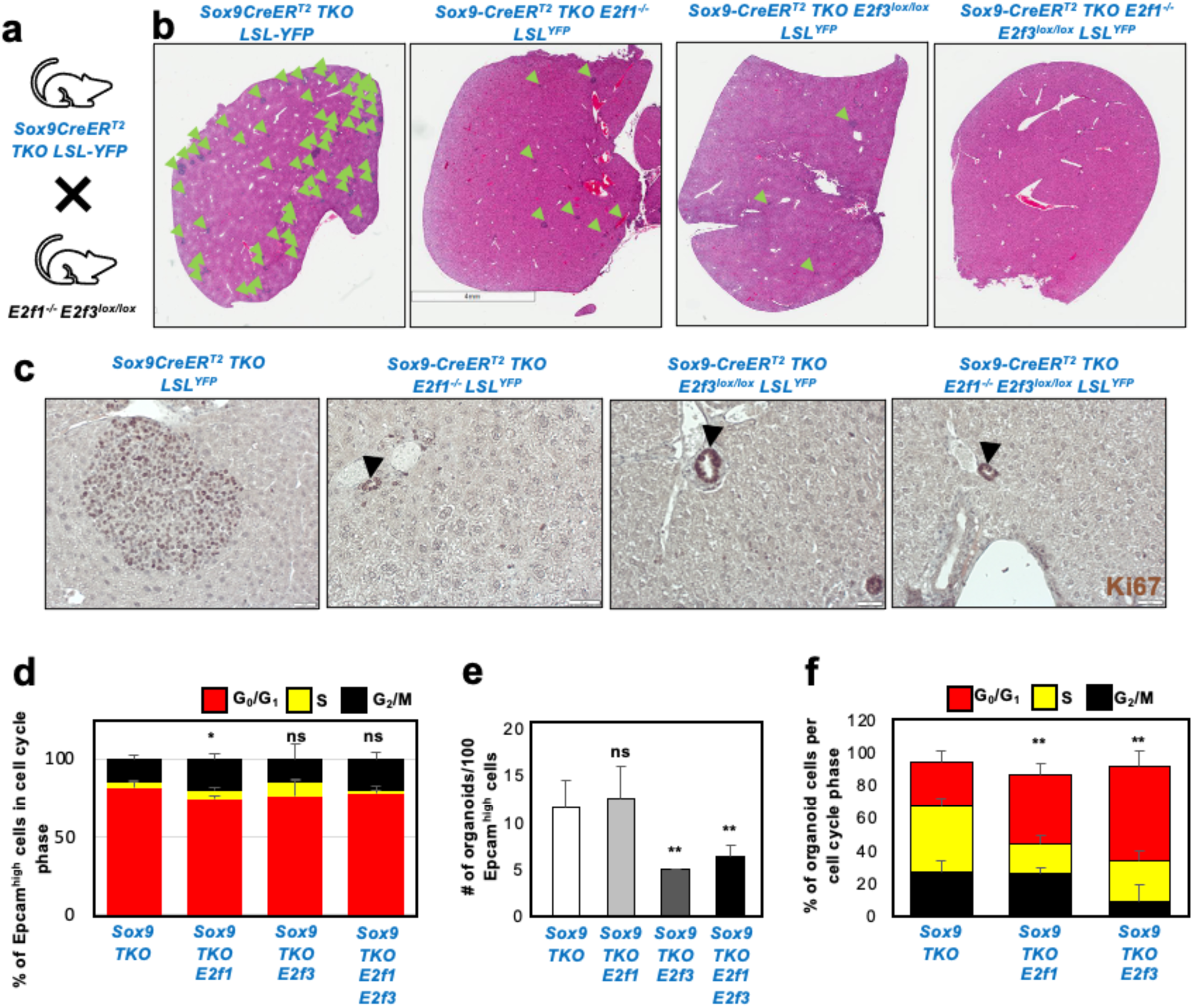
Single and compound *E2f1*/*E2f3* deficiency impairs TKO HCC *^Sox^*^9*-CreERT*2^ initiation without disrupting proliferation. **a**: breeding strategy to incorporate single and double deficiency for *E2f1* and *E2f3* in the c*TKO* background. **b**: Representative H&E staining for one liver lobe from each genotype, twelve weeks after Tamoxifen treatment. Green arrow indicates an individual TKO HCC lesion (n=4 per genotype). **c**: Representative IHC for Ki67 in livers from *Sox9-CreER^T^*^2^ *TKO LSL^YFP^* mice either control, single or double deficient for *E2f1* and *E2f3*. **d**: Cell cycle activity (PI incorporation) of Epcam^high^ cells isolated from the different backgrounds nine weeks after Tamoxifen treatment (n=4). **e**: Organoid forming assay for 100 Epcam^high^ cells isolated from each background (n=5). **f**: Cell cycle activity (PI incorporation) of organoid cells derived from Epcam^high^ cells from each background (n=4).

### Activation of a Sox9/Foxa2 switch by E2f in TKO HCC

Disruption of the transcriptional networks enforcing lineage fidelity tends to be critical for transdifferentiation^33–35^. To determine whether E2f drives the transdifferentiation of Sox9^+^ bile duct cells by altering the integrity of their transcriptional networks, we performed ATAC-Seq analysis of flow-purified Epcam^high^ and Epcam^-^ cells from *Sox9-CreER^T^*^2^ *TKO LSL^YFP^* mice at the nine-week time point. Evolution from Epcam^high^-to Epcam^-^-chromatin states is accompanied by a ∼10% decrease in the number of open chromatin regions (from 44,287 peaks in Epcam^high^ cells to 40,769 peaks in Epcam^-^ cells) (**Fig.5a**; **Fig.S5a-b**). Sequence enrichment analysis shows a decreased contribution of the pioneer factor Sox9 (associated with bile duct identity^27,36^) from Epcam^high^ to Epcam^-^ evolution. In contrast, pioneer factors Foxa2 and Foxa3 (associated with the establishment of hepatocyte identity^37–39^ ^40^) maintain their contribution from Epcam^high^ cells to Epcam^-^ cells evolution (**Fig.5b**). SCENIC^41^ analysis of scRNA-Seq from YFP^+^ cells isolated from *Sox9-CreER^T^*^2^ *TKO LSL^YFP^* mice at the nine-week time point also identifies this inversed pattern of expression and activity between Sox9 and Foxa2/Foxa3 pioneer factors from the “Mix” cluster to the “Hepatocyte” cluster (**Fig.5c**). In addition, IHC staining at different steps of TKO HCC *^Sox^*^9^*^-CreERT^*^2^ development confirms a progressively decreased Sox9 expression, stable Foxa2 expression and increased expression of Foxa3 in a mosaic fashion (**Fig.5d**). Comparative ATAC-Seq analysis of Epcam^high^ cells isolated from the liver of *Sox9-CreER^T^*^2^ *TKO LSL^YFP^* and *Ck19-CreER^T^*^2^ *TKO LSL^YFP^* mice shows that peak identity is largely conserved between both populations, but these peaks tend to be narrower in Ck19^+^-derived Epcam^high^ cells (**Fig.S5c-d**). Foxa3 is not expressed in the hamartoma structure developing in *Ck19-CreER^T^*^2^ *TKO LSL^YFP^* mice (**Fig.S5e**), indirectly suggesting that induction of *Foxa3* expression upon *Rb* family inactivation could represent a key event in the transdifferentiation of Sox9^+^ bile duct cells into tumor cells from the hepatocyte lineage. Supporting this hypothesis, analysis of *Foxa3* expression at the single cell level confirms its progressive increase from the three-week to the five-week time points of TKO HCC development (**Fig.5e-f**).

Since the conformation of *Sox9*, *Foxa2* and *Foxa3* genomic loci is unchanged from Epcam^high^ to Epcam^-^ cells (**Fig.S5f**), we reasoned that their altered expression during TKO HCC development may be due to transcriptional variations rather than epigenetic alterations. Supporting this concept, promoter sequence analysis shows that *Foxa2* and *Foxa3* regulatory regions harbor evolutionary conserved low-affinity *E2f* binding sites^18^ (**Fig.S5g**). ChIP assays confirm that these putative *E2f* binding sites are functional in TKO HCC (**Fig.5g**), identifying *Foxa2* and *Foxa3* as non-canonical *E2f* target genes in the liver. While Sox9 and Foxa2 expression is not affected by *E2f1* and/or E2f3 deficiency, Foxa3 expression decreases upon single and compound *E2f1* and *E2f3* deficiency (**Fig.5h**), indicating that *Foxa3* expression specifically relies on *E2f* activity. Together, these results identify an inversed pattern of activity and expression between Sox9 and Foxa factors during TKO HCC development, which we refer to as the “Sox9/Foxa switch”.

### The Sox9/Foxa switch is a functional feature of TKO HCC

To functionally determine the consequences of the Sox9/Foxa switch for TKO HCC development, we modulated their expression in organoids derived from Epcam^high^ cells isolated from *Sox9-CreER^T^*^2^ *TKO LSL^YFP^* mice. We first found that ectopic *Sox9* expression decreases the organoid forming activity of Epcam^high^ cells (**Fig.6a**). In contrast, *Foxa2* and *Foxa3* silencing decreases the organoid forming activity of Epcam^high^ cells (**Fig.6b-d**) as well as the proliferation of TKO HCC derived cells^14,18^ (**Fig.6e-f**). Moreover, ectopic expression of *Foxa3* in Epcam^high^ cells isolated from *Ck19-CreER^T^*^2^ *TKO LSL^YFP^* mice at the nine-week time point increases their organoid forming activity (**Fig.6g**). Finally, ectopic expression of *Foxa3* in human HCC cell lines represses *Sox9* expression in two lines out of three, suggesting that the repression of *Sox9* expression, as observed in **Fig.5**, may be a direct consequence of increased Foxa3 activity. Collectively, these functional assays show an opposite role for *Sox9* and *Foxa* pioneer factors in TKO HCC development.

**Figure 5.**
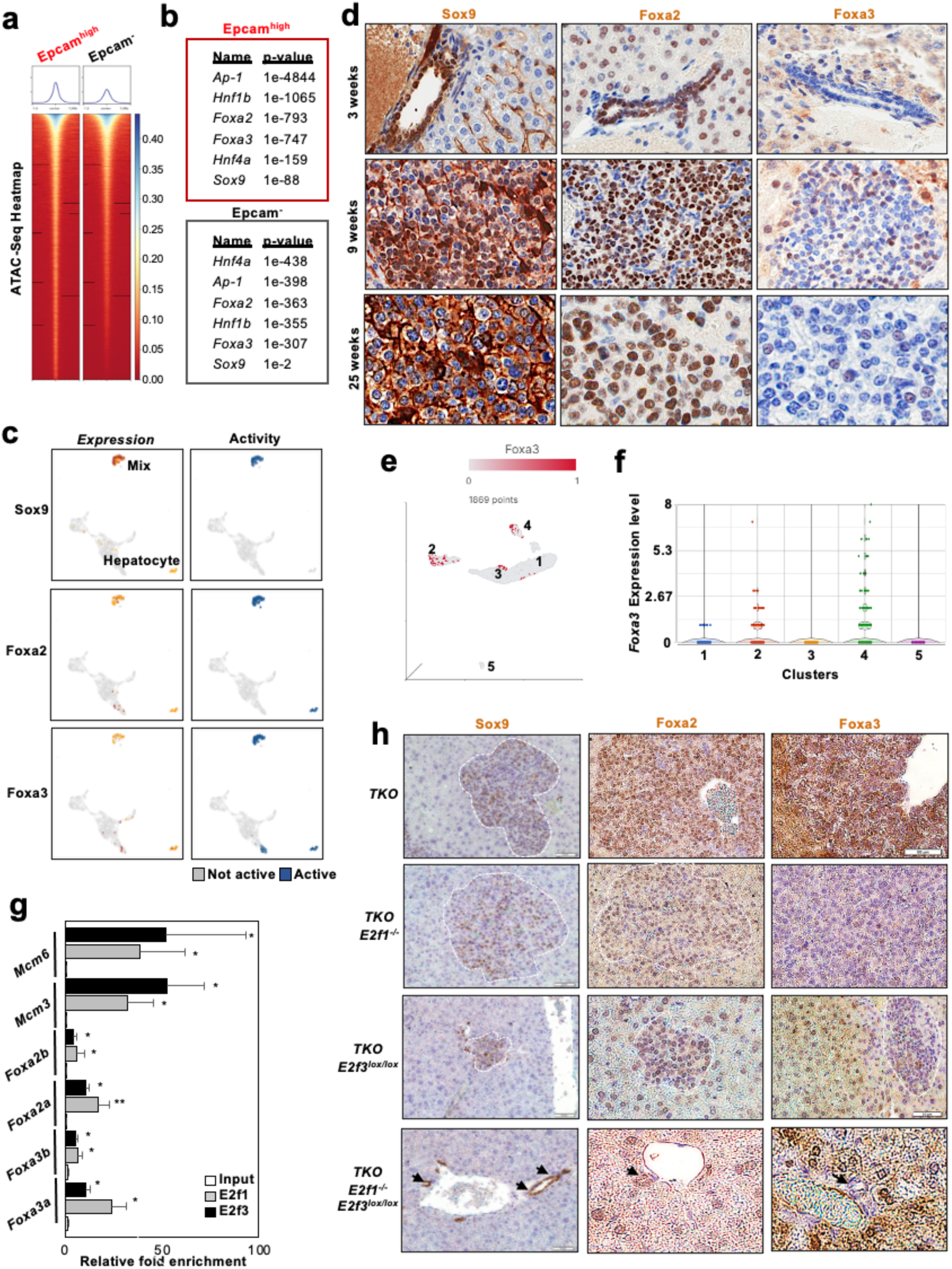
**a**: Heatmap for the ATAC-Seq analysis of Epcam^high^ and Epcam^-^ cells from *Sox9-CreER^T^*^2^ *TKO LSL^YFP^* mice isolated at the nine-week time point. The upper panels indicate the number (y axis) of open regions based on their length (x axis) for each population. **b:** Motif enrichment analysis of ATAC-Seq data identifies the enrichment of relevant transcription factor binding sites in the open chromatin regions for each population. **c**: UMAP display of SCENIC analysis of scRNA-Seq data (as in **Fig.1x**) for Sox9, Foxa2 and Foxa3 expression (left panel; heatmap visualization-red:high; pale yellow:low) and activity (right panel-binary visualization) in TKO HCC*^Sox9CreERT^*^2^. **d:** IHC staining for Sox9, Foxa2 and Foxa3 at the three-, nine- and 25-week time points following Tamoxifen treatment of *Sox9-Cre-ER^T^*^2^ *TKO* mice. *NB*: Sox9 is a nuclear protein and intercellular staining is background signal. **e**: UMAP display of *Foxa3* expression in scRNA-Seq datasets from YFP^+^ cells from *Sox9-CreER^T^*^2^ *TKO LSL^YFP^* at three- and nine-week time points, as displayed in **Fig.2e**. **f**: Violin plot display of *Foxa3* expression in each cluster, as displayed in **Fig.2e**. **g**: ChIP assay for E2f1 and E2f3 in primary TKO HCC tumors shows that E2f binds to *Foxa2* and *Foxa3* promoter regions. *Mcm3* and *Mcm6* promoter regions are used as positive controls (n=3). **h:** Representative IHC for Sox9 (left panels), Foxa2 (middle panels) and Foxa3 (right panels) expression in the livers of *Sox9-CreER^T^*^2^ *TKO LSL-YFP* mice (TKO) with single or compound *E2f1* and *E2f3* deficiency.

**Figure 6:**
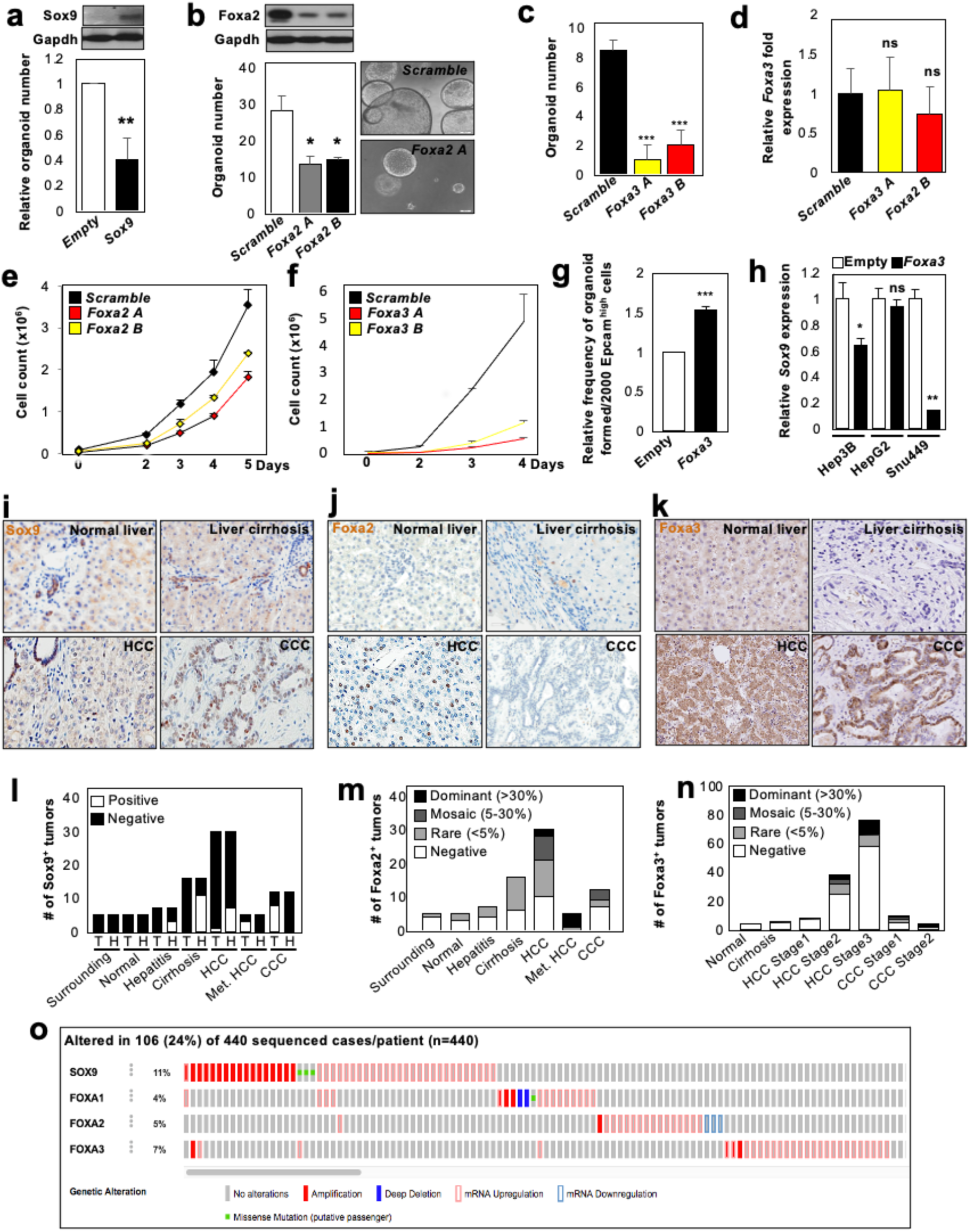
**a**: Upper panel: immunoblotting for Sox9 expression upon its ectopic expression in organoids. Lower panel: relative organoid forming activity of 100 Epcam^high^ cells upon expression of *Sox9* (n=3). **b:** Upper panel: immunoblotting for Foxa2 expression upon its silencing with two distinct hairpins in organoids. Lower left panel: organoid forming activity of 100 Epcam^high^ cells upon repression of *Foxa2* expression by shRNA (n=3). Lower right panel: representative picture of organoids upon repression of *Foxa2* expression by shRNA. **c**: organoid forming activity of 100 Epcam^high^ cells upon repression of *Foxa3* expression by shRNA (n=3). **d**: RT-qPCR detection of *Foxa3* expression in organoids depicted in **Fig.6c** shows that only organoids escaping *Foxa3* silencing have grown (n=3). **e:** Growth curve for a TKO HCC derived cell line upon the expression of *Scramble* control or shRNAs targeting *Foxa2* (hairpin A and B) (n=3). **f**: Growth curve for a TKO HCC derived cell line upon the expression of *Scramble* control or shRNAs targeting *Foxa3* (hairpin A and B) (n=3). **g:** Ectopic expression of *Foxa3* in Epcam^high^ cells freshly isolated from *Ck19-CreER^T^*^2^ *TKO LSL-YFP* mice at the nine week time point increases their organoid forming activity (n=3). **h**: RT-qPCR analysis of *Sox9* expression upon ectopic expression of *Foxa3* in three human HCC cell lines (n=3). **i-k:** Representative IHC for Sox9 (**i**), Foxa2 (**j**) and Foxa3 (**k**) in tumor microarray (TMA) at different stages of human HCC development: normal liver (normal tissue); liver cirrhosis (pre-malignant stages); HCC and CCC (cholangiocarcinoma). **l:** Quantification of Sox9 expression in the different stages of human HCC development. Positivity/negativity was determined by EEF, a board-certified pathologist. **m:** Quantification of Foxa2 expression in the different stages of human HCC development. Stages were divided in subgroups (Rare, Mosaic and Dominant) based on the percentage of Foxa2-expressing cells in each sample, as determined by EEF. Met HCC: metastatic HCC. **n**: Quantification of Foxa3 expression in the different stages of human HCC and CCC development. Stages were divided in subgroups (Rare, Mosaic and Dominant) based on the percentage of Foxa3-expressing cells in each sample, as determined by EEF. **o**: TCGA analysis for genetic alteration targeting *Sox9*, *Foxa1*, *Foxa2* and *Foxa3* in human HCC (440 samples).

To determine the relevance of the *Sox9/Foxa* switch in the context of human HCC, we first performed IHC against Sox9, Foxa2 and Foxa3 in human tissue microarray (TMA) of liver samples from pre-malignant to advanced HCC: Sox9 is expressed in premalignant tissues (hepatitis and cirrhosis) and hamartoma-like structures found at the border of HCC but is largely absent in established HCC (Stage II and onwards) (**Fig.6i, l**). In contrast, Foxa2 expression progressively increases during HCC development and develops a mosaic expression pattern in advanced HCC (**Fig.6j, m**; **Fig.S5h**). Finally, Foxa3 expression also increases during HCC development (**Fig.6k, n**). In addition, TCGA analysis shows that overexpression of *Sox9* and *Foxa* family members is mutually exclusive in human HCC (**Fig.6o**). Interestingly, concomittant *Foxa* gene alterations are not observed in the same sample, suggesting overlapping functions in hHCC. Collectively, these data indicate that the Sox9/Foxa switch is a functional feature of HCC.

### Foxa3 rewires the metabolism of Sox9^+^ bile duct cells to promote TKO HCC initiation

Among *Foxa* factors, *Foxa3* exerts a particularly potent role in liver regeneration and transdifferentiation of fibroblasts into hepatocytes^40,42^. As our data in **Fig.5&6** suggest a critical role for *Foxa3* in TKO HCC development from Sox9^+^ bile duct cells, we sought to identify the molecular mechanisms driven by *Foxa3* to drive the transdifferentiation of Sox9^+^ bile duct cells towards the hepatocyte lineage. Partitioning “Mix” cluster cells (**Fig.2e**) based on *Foxa3* expression, we first found a negative correlation between *Foxa3* and bile duct marker (*Krt19*, *Sox9* and *Epcam*) expression (**Fig.7a; Table S7**). Filtering genes that are positively correlated with *Foxa3* expression with genes that are repressed upon triple inactivation of *Foxa* factors in liver^37^, we generated a list of *Foxa3* target genes in TKO HCC. Ingenuity Pathway Analysis (IPA) of these target genes reveals the enrichment for enzymes driving several metabolic pathways (urea cycle; tyrosine catabolism; beta-oxidation and ketogenesis, both involved in fatty acid metabolism^43^-**Fig.S6a**) that are hallmarks of periportal hepatocytes^44^, suggesting that *Foxa3* alters the cell fate of TKO Sox9*^+^* bile duct cells by rewiring their metabolic activity. We tested this hypothesis by experimentally focusing on beta oxidation and its downstream ketogenesis pathways since beta oxidation can be pharmacologically targeted by Trimetazidine® (TMZ®)^45^. Treatment of *Sox9-CreER^T^*^2^ *TKO LSL^YFP^* mice with TMZ® leads to a ∼two-fold reduction of the tumor burden (**Fig.7b-c**), similarly affecting both Epcam^high^ and Epcam^-^ subpopulation frequency (**Fig.7d**). In contrast, a similar regimen does not significantly reduce the tumor burden in *Alb-CreER^T^*^2^ *TKO LSL^YFP^* mice (**Fig.7b-c**). However, while TMZ® treatment in *Sox9-CreER^T^*^2^ *TKO LSL^YFP^* mice decreases tumor cell proliferation (**Fig.7e**), it triggers apoptotic activity in *Alb-CreER^T^*^2^ *TKO LSL^YFP^* mice (**Fig.7f**), identifying distinct responses of TKO HCC *^Sox^*^9^*^-CreERT^*^2^ and TKO HCC *^Alb-CreERT^*^2^ to TMZ® treatment. Finally, comparative scRNA-Seq analysis of CT and TMZ® treated YFP^+^ cells show that TMZ® treated cells are spatially distinct within the same cluster (**Fig.7g; Table S8**). In particular, TMZ® treatment decreases Ki67 and Hnf4a expression in “Mix” cluster cells (**Fig.7h**), supporting the concept that activation of beta oxidation by Foxa3 promotes the transdifferentiation of Sox9^+^ bile duct cells into the hepatocyte lineage.

Collectively, our results identify Sox9^+^ bile duct cells as a cell of origin for HCC upon disruption of the Rb/E2f disruption, a hallmark of HCC. In addition to the activation of a canonical cell cycle program, our data reveal that E2f transactivates the pioneer factor *Foxa3* to trigger the transdifferentiation of Sox9^+^ bile duct cells into periportal hepatocyte-like cells and initiate a HCC that displays a striking phenotypic overlap with HCC developing from periportal hepatocytes in the same genetic context (**Fig.S6b**).

## DISCUSSION

### A novel model for the role of E2f in cancer initiation

Disruption of Rb/E2f interaction is a hallmark of cancer initiation. In the current model, the resulting E2f oncogenic activity (i.e. unrestricted proliferation) is an extension of its physiological role ^18,31,46,47^. Departing from this paradigm, we believe that our data support a more complex role for E2f in cancer. In this new model, we propose that the disruption of the Rb/E2f interaction generates two distinct and non-linear consequences: *i)* the first consequence is the de-repression of cell cycle genes, which triggers their transactivation by several transcription factors such as E2f^32^, but also Myc^32^, Foxm1, Bmyb^48^, etc, overall leading to sustained proliferation. *ii)* the second consequence is the activation of non-cell cycle oncogenic features by E2f in a cellular specific context. In particular, we previously showed that E2f target genes can be partitioned into two categories based on E2f binding affinity for their regulatory regions^18^: while high affinity target genes are promptly activated by E2f upon acute disruption of the Rb/E2f interaction, low affinity target genes, such as *Foxa3*, are only transactivated in the context of sustained disruption of the Rb/E2f interaction. Therefore, we propose that the specific contribution of E2f to cancer resides in the context-dependent acquisition of novel non-cell cycle functions rather than the extension of its physiological cell cycle functions. From a translational point of view, this model suggests that targeting these non-cell cycle features could represent an alternative therapeutic strategy in the context of resistance to CDK inhibitors. In addition, our results also suggest that the individual targeting of activator E2f1 and E2f3 could significantly repress E2f oncogenic activity. In this context, we previously showed that their interactome are completely distinct, suggesting that the unique oncogenic contribution of individual E2f factors to cancer, and therefore their therapeutic vulnerability, may reside in the recruitment of co-factors.

### A ductal origin for HCC

Mature hepatocytes are generally considered as the only cell of origin for HCC, largely based on carcinogen-induced models of HCC ^12,29^. However, these models generate an inherent bias since carcinogen activation relies on *Cyp2e1*, which is predominantly expressed in pericentral hepatocytes^49^. Therefore, whether other liver populations can serve as a cell of origin for HCC remains unclear, especially since these populations are also exposed to the main HCC risk factors. Our data show that both periportal hepatocytes and Sox9^+^ bile duct cells can serve as a cell of origin for a phenotypically similar HCC upon disruption of the Rb/E2f interaction. However, the kinetic of HCC development from Sox9^+^ bile duct cells is much slower than its counterpart initiating from periportal hepatocytes. Reasons for this kinetic difference are probably multiple, such as the fact that the transdifferentiation from bile duct to hepatocyte lineage is inherently inefficient, or because Sox9^+^ bile duct cells are less abundant than periportal hepatocytes and/or display a lower mitotic index. In experimental contexts where both populations simultaneously initiate HCC, we postulate that this competitive disadvantage either prevents HCC *^Sox9+^* cells from even expanding or force their blending into more abundant HCC *^Alb+^* tumor cells, especially since they share common histological features. From a translational perspective, the fact that HCC *^Alb-CreERT^*^2^ and HCC *^Sox^*^9^*^-CreERT^*^2^ display remarkably similar molecular and histological features at advanced stage points suggests that the ductal origin of HCC originating from bile duct cells can be difficult to establish when diagnosed at an advanced stage, as often the case in the clinic. However, our data suggest that discrete markers could help classify HCC based on their ductal or hepatocyte origin (**Fig.1**), which could prove useful as HCC initiated from bile duct cells appears to retain unique therapeutic susceptibilities related to their origins (**Fig.7**).

**Figure 7.**
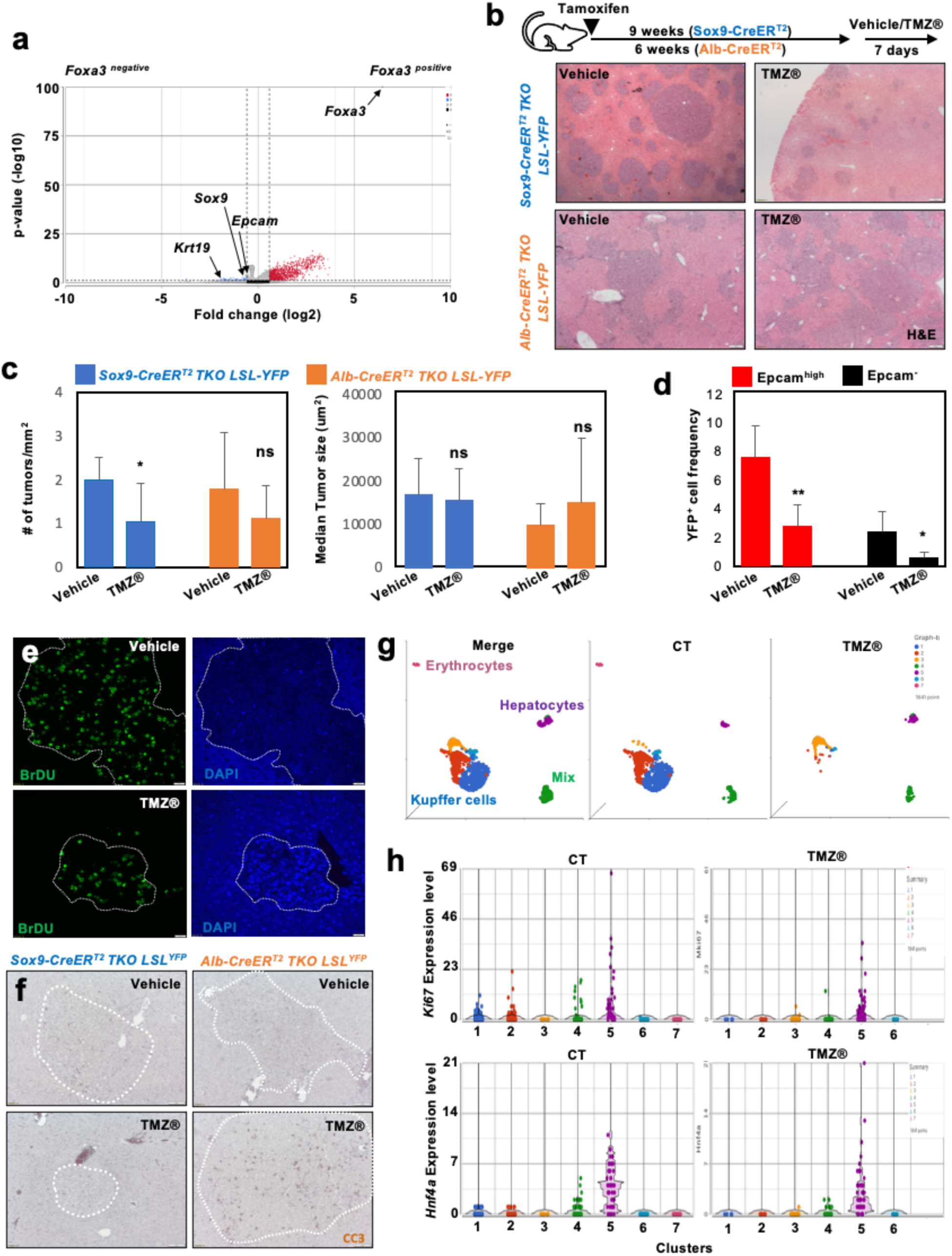
**a:** Cells from clusters 2 and 4, as displayed in **Fig.2e**, were partitioned based on *Foxa3* expression. Comparative gene analysis identifies gene differentially expressed between *Foxa3* positive and *Foxa3* negative cells. **b**: Upper panel: Experimental protocol to treat *Sox9-CreER^T^*^2^ *TKO LSL^YFP^* (9 weeks post Tamoxifen treatment; n=6/group) and *Alb-CreER^T^*^2^ *TKO LSL^YFP^* (6 weeks post Tamoxifen treatment; n=5/group) mice with either vehicle or Trimetazidin® (TMZ®) for 7 days. Lower panels: representative H&E staining of liver from both lines and regimens. **c**: quantification of the number of tumors per mm^2^ (left panel) and median tumor size (right panel) in mice from the different groups, as depicted in **b**. **d**: *Sox9-CreER^T^*^2^ *TKO LSL^YFP^*were treated with either vehicle or TMZ® for 7 days, 9 weeks post Tamoxifen treatment. Livers were digested to quantify the frequency of Epcam^high^ YFP^+^ and Epcam^-^ YFP^+^ cells in each group (n=7/group). **e**: Representative co-immunofluorescence for BrdU in *Sox9-CreER^T^*^2^ *TKO LSL^YFP^* mice treated with either vehicle or TMZ® for one week, 9 weeks post Tamoxifen treatment (as in **b**). Slides were counterstained with DAPI. **f**: Representative IHC staining for cleaved caspase 3 (CC3) in *Sox9-CreER^T^*^2^ *TKO LSL^YFP^* and *Alb-CreER^T^*^2^ *TKO LSL^YFP^* treated with either vehicle or TMZ® for one week (as in **b**). **g**: UMAP display of merged scRNA-Seq datasets from YFP^+^ cells isolated from *Sox9-CreER^T^*^2^ *TKO LSL^YFP^* at 9-week time point, either control (CT) or treated with TMZ® for 7 days. Left: merge display; clusters are identified based on marker expression; Center and Right: individual displays. **h**: Violin plot display of *Ki67* (upper panel) and *Hnf4a* (lower panel) expression in each cluster, as displayed in **Fig.7g**.

## MATERIAL AND METHODS

### Mice

*Rb*-family *cTKO* mice were bred as previously described^31^. *Hnf4a-Cre* (EMMA repository #02506), *Foxa3-Cre*^23^, *Foxl1-Cre*^50^, *Gja1-CreER^T^*^2^ (EMMA repository #00330), *Ck19-CreER^T^*^2^ ^22^ and *Sox9-CreER^T^*^2^ ^51^ mice were previously described. For *Hnf4a-Cre* and *Foxa3-Cre TKO* mice, the germline loss of *Rb* family members impairs viability at 6-8 week-time point. Long-term observations are performed on *Rb^lox/lox^*, *p130^lox/lox^*, *p107^+/-^*(defined as *p107* single) and displayed in **Fig.1**. *Rosa26^LSL-YFP^* mice were obtained from the Jackson Laboratory (*Gt(ROSA)26Sortm1(EYFP)Cos/J*). E2f1^-/-^ E2f3^lox/lox^ mice are a generous gift of Dr. Gustavo Leone. All experiments with mice were approved by CHOP IACUC (protocol #969). *Subcutaneous transplantation:* organoids were physically disrupted and mixed with Matrigel prior to transplanting in the flank of NSG mice. TMZ® treatment: Trimetazidine® was resuspended in a mix of DMSO/PEG300/Tween80/water to deliver a daily dose of 100mgr/kg by gavage.

### Cell Culture

*Rb*-family TKO 1.1 and 2.1 cell lines were derived from independent TKO HCC tumors and have been previously described^30^. Experiments have been performed in both cell lines with similar results. TKO1.1 and 2.1 cell lines are collectively referred to as TKO HCC cells. TKO HCC, HepG2 and Hep3B cells were cultured in DMEM (Mediatech), supplemented with 10% FBS/ 1% Penicillin-Streptomycin-Glutamine (Invitrogen). SNU449 cells were cultured in RPMI, supplemented with 10% FBS/ 1% Penicillin-Streptomycin-Glutamine. Organoid cultures were performed as described^52^. Single cell were resuspended in Matrigel and plated in drops of 50ul for growth assay (30ul for pharmacological inhibition of Yap signaling). *Viral infection:* shRNA sequences were generated using psicoligomaker^64–66^ and cloned into the pSicoR-PGK-RFP and psicoR-CMV-GFP vectors. Foxa3 cDNA was cloned into pUltra-GFP. Viral infection was performed as previously described and infected cells were isolated 2-4 days post infection on the basis of GFP/RFP expression by flow cytometry^31^.

### In silico RNA-Seq, ATAC-Seq, Fluidigm assay, 10x genomics

#### Sample preparation

<u>RNA-Seq</u>: cells isolated by flow on the basis of Epcam expression were directly sorted into Trizol LS. Trizol-extracted RNA was processed for bulk RNA-Seq by the Next Generation Sequencing Core at Penn Medicine. <u>ATAC-Seq</u>: ATAC-seq libraries were prepared as described^53^. In brief, 50,000 cells were isolated by flow cytometry and washed with cold 1x PBS. Cell pellet was resuspended in 50 µl of cold lysis buffer (10 mM Tris-HCl, pH 7.4, 10 mM NaCl, 3 mM MgCl_2_ and 0.1% (v/v) Igepal CA-630) and immediately spun at 1,600x*g*, 4 °C for 10 min. Nuclei pellet was resuspended in 50 µl of transposition reaction mix (1x Tagment DNA Buffer, 2.5 µl of Tagment DNA Enzyme 1 (Illumina)) and incubated at 37 °C for 30 min. Subsequent steps of the protocol were performed as previously described (Buenrostro et al., 2013). The libraries were purified using a Qiagen MinElute Gel Purification Kit and library concentration was measured using both Qubit and KAPA qPCR. 2100 Bioanalyzer was used to check the quality of the libraries. The libraries were sequenced on the Illumina NextSeq 500, with 75-bp paired-end reads. <u>10x genomics</u>: to avoid any bias in the selection of cells to be processed for scRNA-Seq analysis, unfractioned YFP^+^ cells were sorted by flow cytometry and resuspended at a concentration of 1000 cells/ul. Cells in suspension were processed by the Next Generation Sequencing Core at Penn Medicine and CHOP Single Cell Biology Core. Of note, this unbiased selection based on YFP expression leads to the presence of autofluorescent Kupffer cells (positive into all channels in the absence of fluorochome at their surface) in analyzed samples, which were identified based on the expression of multiple key markers.

#### Data processing

<u>ATAC-seq</u>: after trimming the adapters with attack (version 0.1.5 – https://atactk.readthedocs.io/en/latest/index.html), the raw reads were aligned to the mm9 genome using bow-tie-1.1.2 ((https://genomebiology.biomedcentral.com/articles/10.1186/gb-2009-10-3-r25) with the following flags: --chunkmbs2000—sam—best—strata-ml-X2000. We used MACS2^54^ for peak calling with a q cutoff of 0.05. Downstream analysis and visualization was performed using HOMER^55^ and deepTools2^56^. <u>scRNA-seq and bulk RNA-Seq</u>: all analysis were performed with Partek (Illumina) and Ingenuity Pathway Analysis (Quiagen). Finally, we reconstructed gene regulatory using the SCENIC (v 1.1.2-2) package^41^.

### qPCR, immunoblotting, staining, flow cytometry

#### qRT-PCR

Total RNA was extracted, purified and reverse transcribed as previously described. qPCR was performed with SYBR Green PCR Master Mix (Life Technologies) on the Viia7 Real-Time qPCR system (Life Technologies). Data were normalized using *gapdh* as a reference gene. Primer sequences are available upon request. *Immunoblotting:* Frozen tumors, control livers, and cells in culture were lysed in 1% SDS or RIPA. *Immunohistochemistry& immunostaining:* Standard deparaffinization, rehydration, and heat-induced epitope retrieval were performed. *Cell Cycle Analysis*: Cell cycle analysis by PI incorporation was performed as previously described^20^. *Antibodies*: Foxa2 (Abcam#108422), Foxa3 (Sigma SAB2108468), Sox9 (Millipore Ab5535), Epcam (ebioscience, clone G8.8 for flow cytometry and Abcam #71916 for IF), Albumin (Bethyl lab A90-234A), Ck19 (a generous gift of Dr. Ben Stanger)^57^, YFP (Abcam#13970), Ki67 (BD#550609), BrdU (BD347580). Human HCC tissue array (LV8011a) was purchased from US Biomax. HCC stages were classified according to the AJCC Clinical Staging System (v7). Analysis of the IHC staining of the tissue array was performed by EEF, a board-certified pathologist. *Flow cytometry*: analysis was performed on a BD Fortessa and BD Canto while sorting was performed on a BD Facs Aria. The surface marker screen on Epcam^high^ cells was performed with the LegendScreen Mouse-PE Kit from Biolegend.

## ACKNOWLEDGMENTS

We thank the Stanger lab for generously sharing their anti-Ck19 antibody. We thank Gustavo Leone for sharing his E2f1/E2f3 deficient mice. We thank the members of the Animal Facility at LIMR as well as the CHOP Single Cell Biology Core. P.V. is supported by American Cancer Society grants (IRG-78-002-35 and RSG-16-233-01-TBE) and RO1 grant (CA251397) from NCI.

## AUTHOR CONTRIBUTIONS

GC and EK designed, performed, analyzed experiments and co-wrote the manuscript; GC and KH performed bioinformatic analysis; EEF supervised and analyzed all pathology-related assessments; PV designed, performed and analyzed experiments, secured funding and supervised the project; PV wrote the manuscript.

---

**Figure S1.**
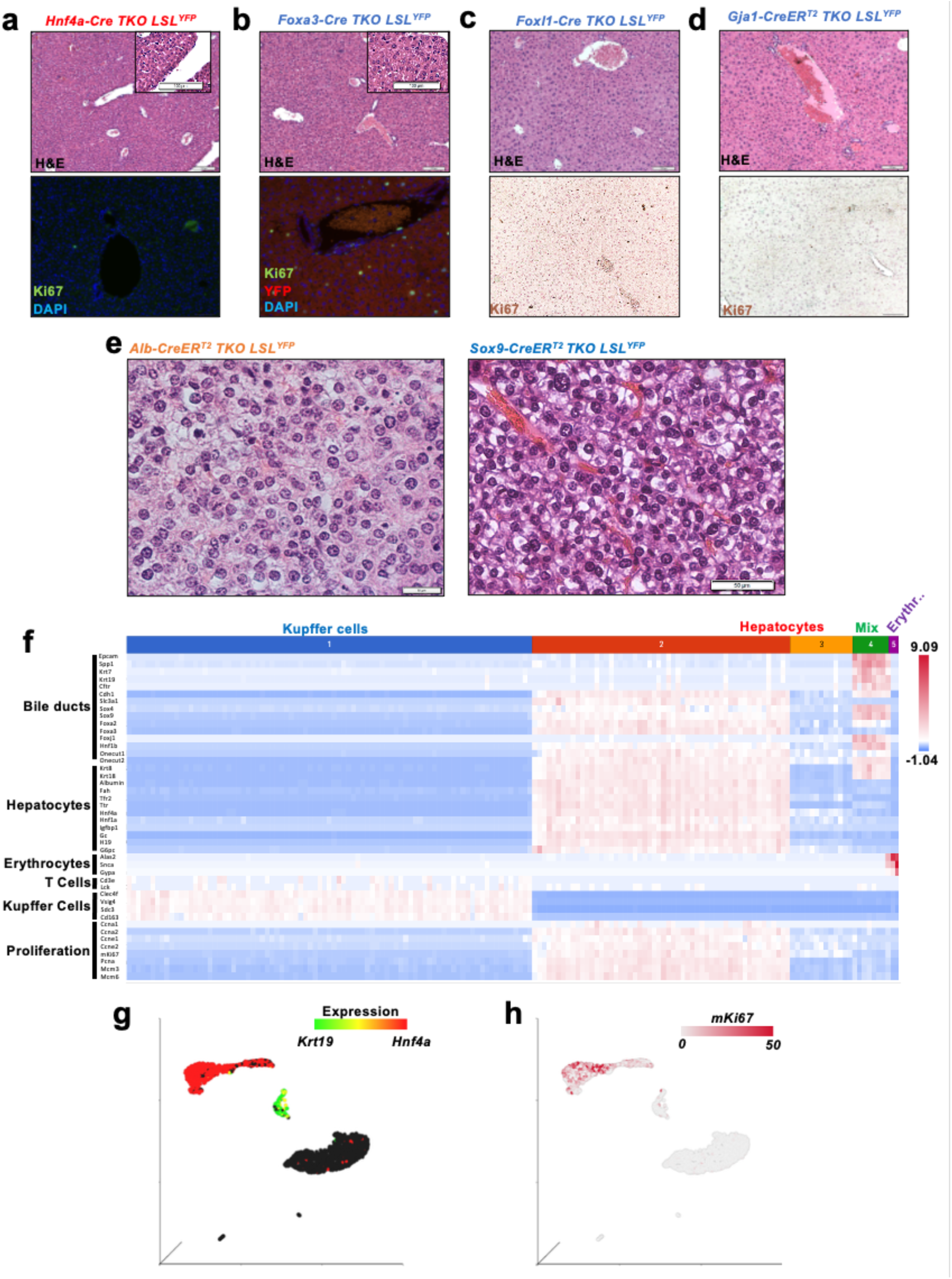
**a:** *Hnf4a-Cre TKO LSL^YFP^* mice do not survive more than 6-8 weeks after birth. Representative H&E staining of a 6-week-old individual shows the absence of a tumor phenotype (upper panel). IF staining for Ki67, counterstained by DAPI, shows little proliferation in these mice at that time point (lower panel). **b:** *Foxa3-Cre TKO LSL^YFP^* mice do not survive more than 6-8 weeks after birth. Representative H&E staining of a 6-week-old individual shows the absence of a tumor phenotype (upper panel). Co-IF staining for Ki67 and YFP, counterstained by DAPI, shows occasional ectopic proliferation in hepatocytes (lower panel), as observed in *Foxa3-Cre p107-single LSL^YFP^* mice .**c:** *Foxl1-Cre TKO LSL^YFP^* are viable and fertile. They display increased frequency of mitotic figures, as evidenced by H&E (up) and Ki67 IHC (down) staining, but no tumor phenotype. **d:** *Gja1-CreER^T^*^2^ *TKO LSL^YFP^*mice do not survive more than two weeks post Tamoxifen treatment. Within this time frame, they do not develop any proliferative phenotype in the liver, as evidenced by H&E (up) and Ki67 IHC (down) staining. **e**: representative high magnification pictures of H&E-stained HCC from *Alb-CreER^T^*^2^ *TKO LSL^YFP^* and *Sox9-CreER^T^*^2^ *TKO LSL^YFP^* show similar histological characteristics. **f**: heatmap showing key lineage and proliferation marker expression in merged scSeq-RNA datasets from *Alb-CreER^T^*^2^ *TKO LSL^YFP^*and *Sox9-CreER^T^*^2^ *TKO LSL^YFP^*, as displayed in **Fig.1x**. **g**: UMAP display of *Krt19* (green) and *Hnf4a* (red) expression in merged scRNA-Seq datasets of YFP^+^ cells from *Alb-CreER^T^*^2^ *TKO LSL^YFP^* and *Sox9-CreER^T^*^2^ *TKO LSL^YFP^*. **h**: UMAP display of *Ki67* expression in merged scRNA-Seq datasets of YFP^+^ cells from *Alb-CreER^T^*^2^ *TKO LSL^YFP^* and *Sox9-CreER^T^*^2^ *TKO LSL^YFP^*.

**Figure S2.**
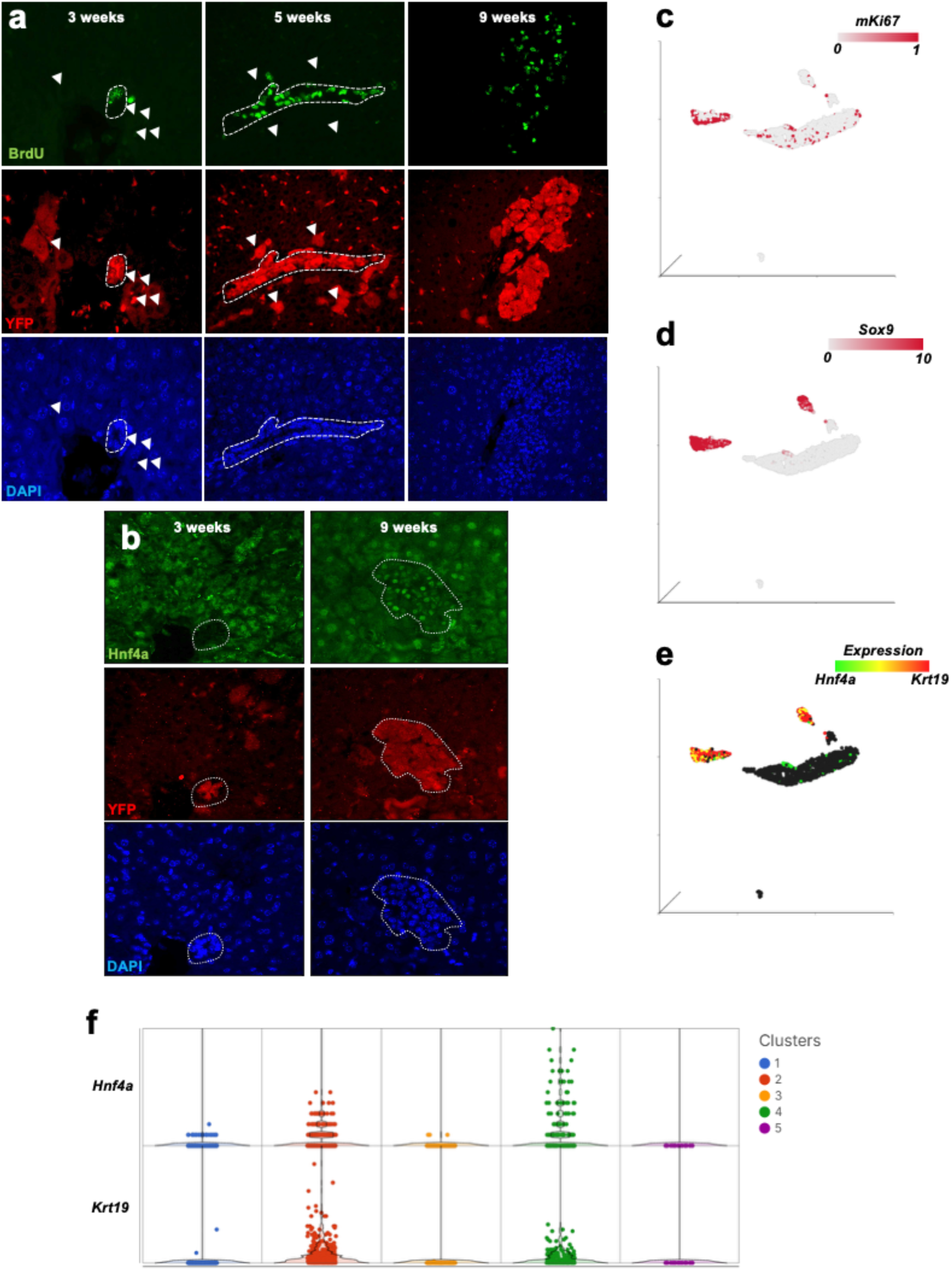
**a:** Co-immunofluorescence staining of livers from *Sox9-CreER^T^*^2^ *TKO LSL^YFP^* mice at three-, five- and nine-weeks post Tamoxifen treatment for BrdU and YFP, counterstained with DAPI. Dotted circles indicate bile ducts (three weeks) and expanding lesions; arrowheads indicate recombined hepatocytes. **b**: Co-immunofluorescence staining of livers from *Sox9-CreER^T^*^2^ *TKO LSL^YFP^*mice at three- and nine-weeks post Tamoxifen treatment for Hnf4a and YFP, counterstained with DAPI. Dotted circles indicate bile ducts (three weeks) and expanding lesions (nine weeks). **c**: UMAP display of *Ki67* expression in merged scRNA-Seq datasets from YFP^+^ cells from *Sox9-CreER^T^*^2^ *TKO LSL^YFP^* at three- and nine-week time points, as displayed in **Fig.2e**. **d**: UMAP display of *Sox9* expression in merged scRNA-Seq datasets from YFP^+^ cells from *Sox9-CreER^T^*^2^ *TKO LSL^YFP^* at three- and nine-week time points, as displayed in **Fig.2e. e**: UMAP display of Krt19 (red) and Hnf4a (green) expression in merged scRNA-Seq datasets from YFP^+^ cells from *Sox9-CreER^T^*^2^ *TKO LSL^YFP^* at three- and nine-week time points, as displayed in **Fig.2e**. **f**: Violin plot display of *Hnf4a* (upper panel) and *Krt19* (lower panel) expression in each cluster, as displayed in **Fig.2e**.

**Figure S3.**
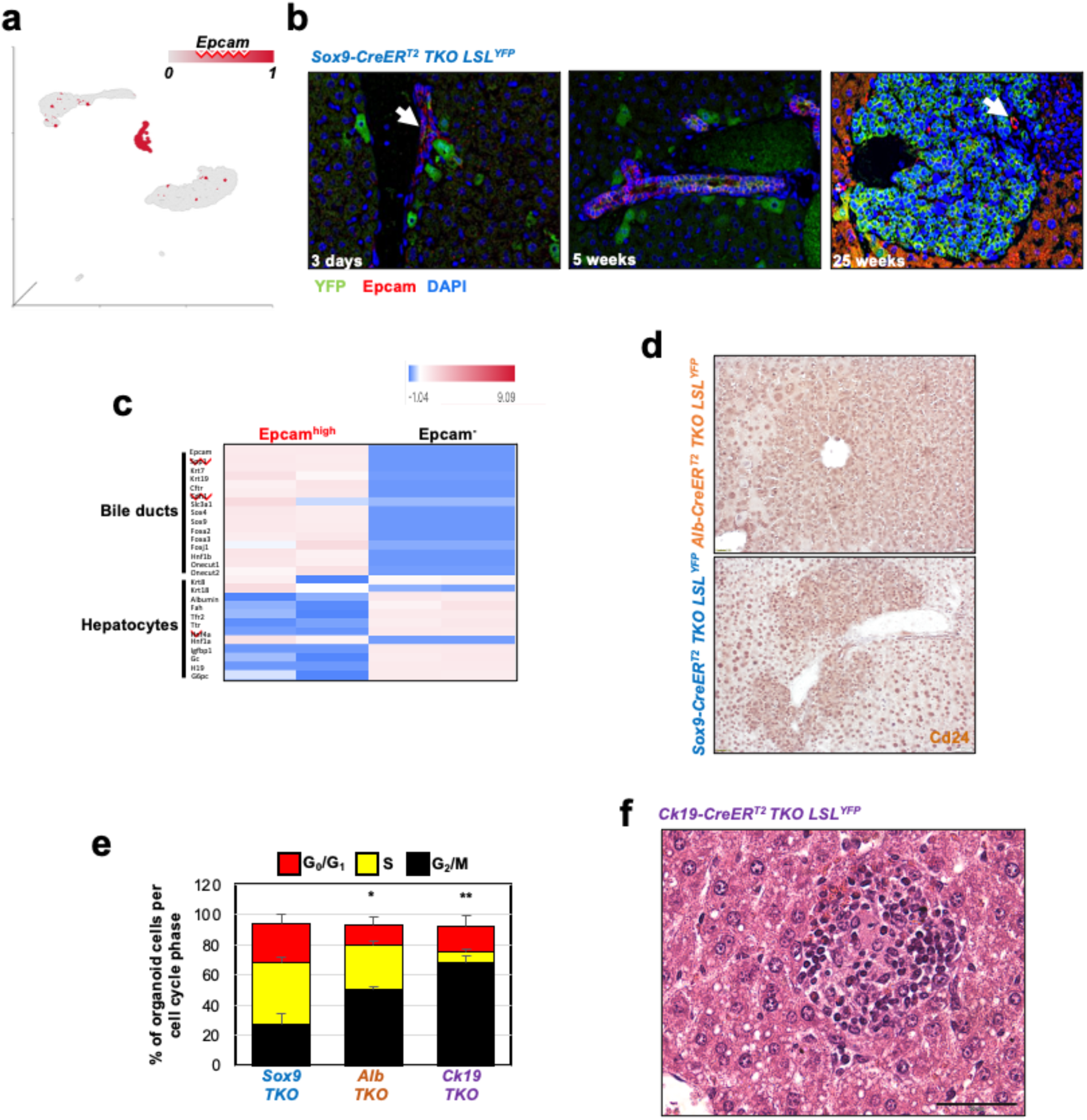
**a:** UMAP display of *Epcam* expression in merged scRNA-Seq datasets of YFP^+^ cells from *Alb-CreER^T^*^2^ *TKO LSL^YFP^* and *Sox9-CreER^T^*^2^ *TKO LSL^YFP^*, as displayed in **Fig.1x**. **b**: Co-IF staining of liver sections from *Sox9-CreER^T^*^2^ *TKO LSL-YFP* mice at three days, five and 25-week time points for Epcam and YFP, counterstained for DAPI. Only merged staining is displayed. **c**: Heatmap for the expression of bile duct and hepatocyte markers (same as in Fig.S1f) in Epcam^high^ and Epcam^-^ cells from TKO HCC (n=2/condition). **d**: Representative IHC staining for Cd24 shows that tumor cells from both *Alb-CreER^T^*^2^ *TKO LSL^YFP^* and *Sox9-CreER^T^*^2^ *TKO LSL^YFP^* express Cd24. **e**: Cell cycle analysis (PI incorporation) of organoid cells from the different lines, as displayed in **Fig.3g**. **f**: A rare TKO HCC early lesion is identified in the H&E staining from a *Ck19-CreER^T^*^2^ *TKO* liver (n=15 different individuals were screened for the presence of TKO HCC lesions).

**Figure S4.**
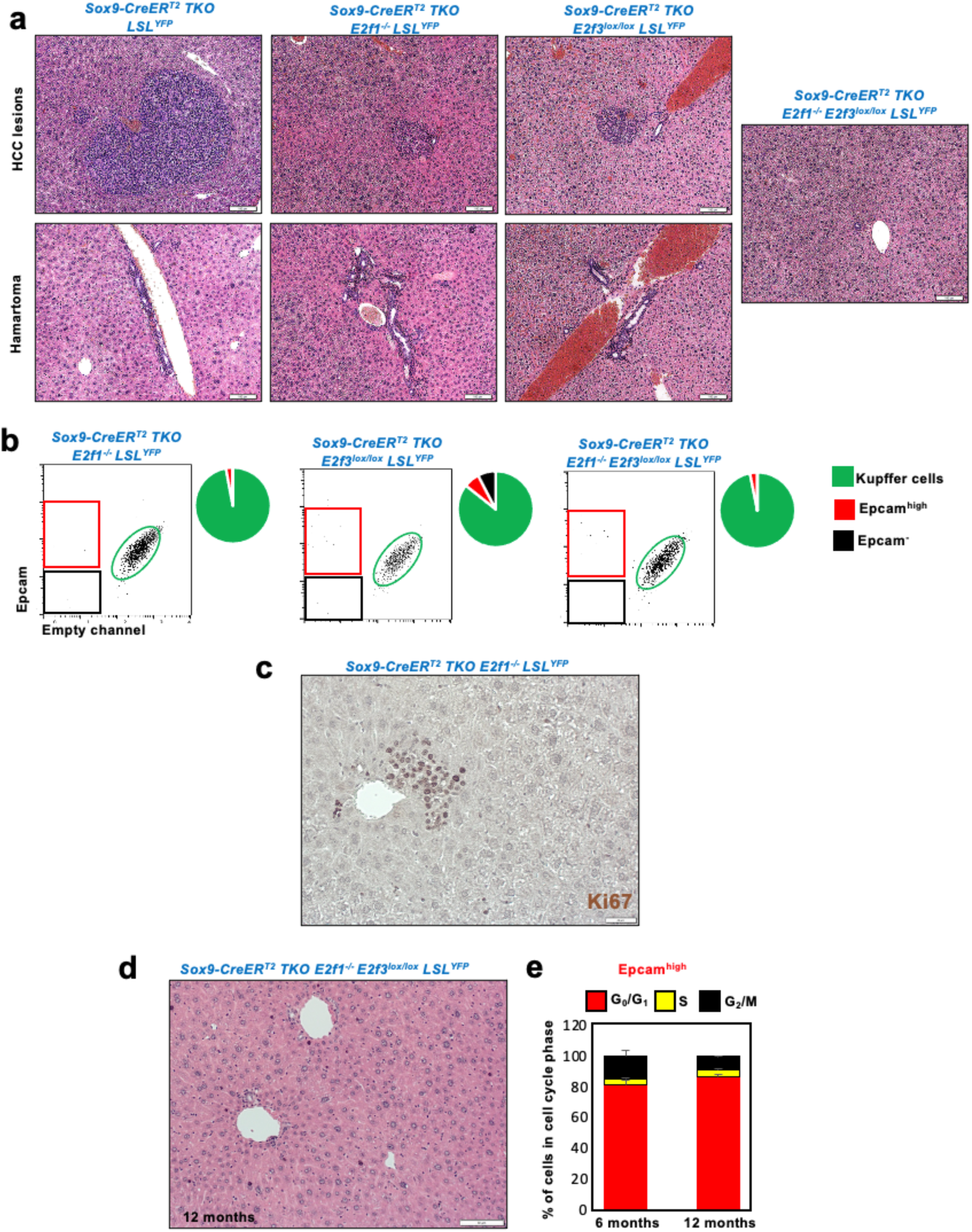
**a:** Representative H&E staining of TKO HCC lesions and hamartomas in the liver of *Sox9-CreER^T^*^2^ *TKO LSL^YFP^* (*TKO*), *Sox9-CreER^T^*^2^ *TKO LSL^YFP^ E2f1^-/-^* (*TKO E2f1^-/-^*), *Sox9-CreER^T^*^2^ *TKO LSL^YFP^ E2f3^lox/lox^* (*TKO E2f3^lox/lox^*) and *Sox9-CreER^T^*^2^ *TKO LSL^YFP^ E2f1^-/-^*, *E2f3^lox/lox^* (*TKO E2f1^-/-^ E2f3^lox/lox^*) mice twelve weeks after Tamoxifen treatment. No HCC lesions or hamartoma can be observed in the latter genotype (n=4). **b**: Frequency of each population, as determined by flow analysis for Epcam in the YFP^+^ fraction from each genotype (n=4). **c**: Representative IHC for Ki67 in TKO HCC lesions developing in *Sox9-CreER^T^*^2^ *TKO E2f1^-/-^ LSL^YFP^* mice. **d**: Representative H&E staining of a liver section from *Sox9-CreER^T^*^2^ *TKO LSL^YFP^ E2f1^-/-^*, *E2f3^lox/lox^* mice 12 months post Tamoxifen treatment (n=10). **e**: Cell cycle analysis (PI incorporation) of Epcam^high^ cells isolated from *Sox9-CreER^T^*^2^ *TKO LSL^YFP^ E2f1^-/-^*, *E2f3^lox/lox^* mice 6- and 12-months post Tamoxifen treatment (n=4 per time point).

**Figure S5.**
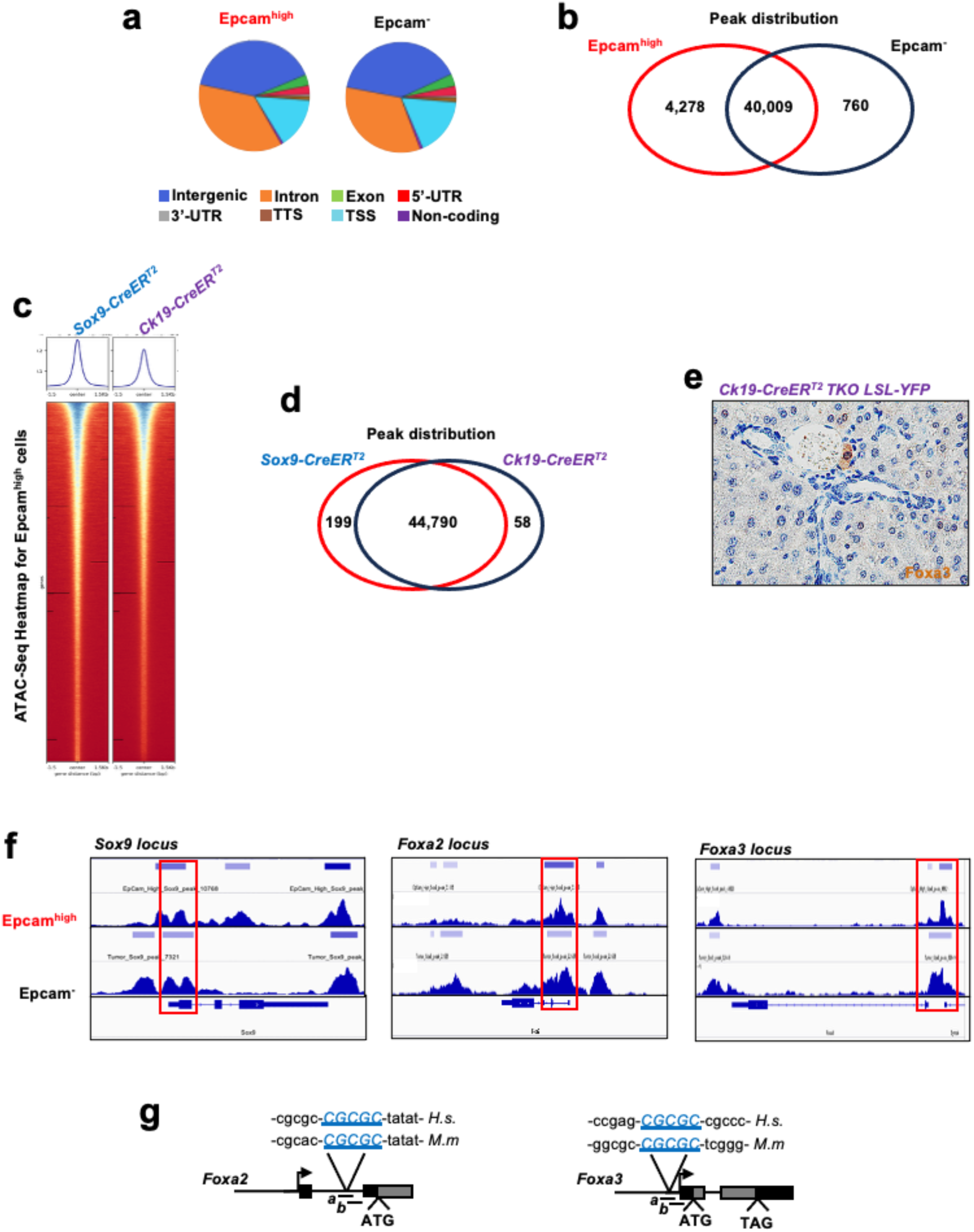
**a:** Open chromatin regions (peaks) in each population, based on their genomic location. **b**: Venn diagram representing the number of peaks in each population. **c**: Heatmap for the ATAC-Seq analysis of Epcam^high^ isolated from *Sox9-CreER^T^*^2^ *TKO LSL^YFP^*and *Ck19-CreER^T^*^2^ *TKO LSL^YFP^* mice at the nine-week time point. The upper panels indicate the number (y axis) of open regions based on their length (x axis) for each population. **d**: Venn diagram representing the number of peaks in each population. **e**: Representative IHC staining for Foxa3 in *Ck19-CreER^T^*^2^ *TKO LSL^YFP^* liver shows that hamartoma structures do not express Foxa3. **f**: Open/close conformation of *Sox9*, *Foxa2* and *Foxa3* locus in Epcam^high^ and Epcam^-^ cells isolated from *Sox9-CreER^T^*^2^ *TKO LSL^YFP^* mice at t^h^e nine-week time point, as determined by ATAC-Seq. **g**: Identification of evolutionary conserved (human/mouse) low-affinity E2f binding sites in the regulatory region of *Foxa2* and *Foxa3*. *a* and *b* represent amplicons used for each gene to amplify the region surrounding the putative E2f binding sites by RT-qPCR in ChIP extracts. *H.s*: Homo Sapiens. *M.m*: Mus Musculus.

**Figure S6.**
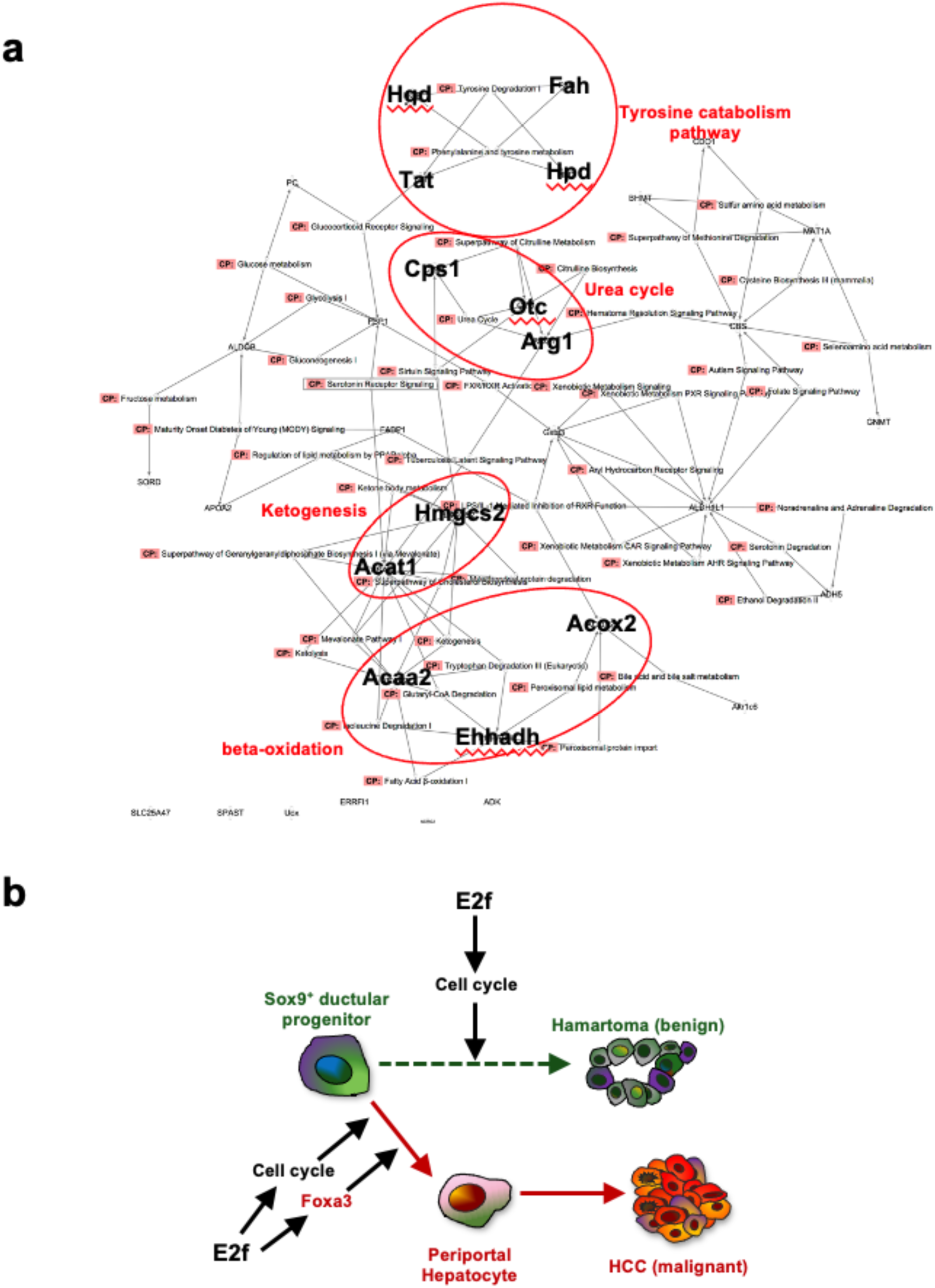
**a**: Ingenuity Pathway Analysis of *Foxa3* target genes in TKO HCC*^Sox^*^9*-CreERT*2^ identifies several metabolic pathways that are specific to periportal hepatocytes. **b**: Model: activation of proliferation in Sox9^+^ bile duct cells leads to hamartoma (a benign bile duct tumor) development, while combined activation of proliferation and *Foxa3* expression promotes their transdifferentiation into periportal hepatocyte-like cells and HCC initiation.

**Table S1, related to Fig.1w**: genes differentially expressed between TKO HCC Sox9-Cre and TKO HCC Alb-Cre

**Table S2, related to Fig.1x**: biomarkers for each cluster

**Table S3, related to Fig.2e**: biomarkers for each cluster

**Table S4, related to Fig.3b**: genes differentially expressed between Epcamhigh and Epcam-cells

**Table S5: related to Fig.3h**: genes differentially expressed between Ck19-derived and Sox9-derived organoids

**Table S6: related to Fig.3i**: genes differentially expressed between Alb-derived and Sox9-derived organoids

**Table S7: related to Fig.7a**: Foxa3 target genes in TKO HCC

**Table S8: related to Fig.7g**: biomarkers for each cluster

